# Genetic analyses led to the discovery of a super-active mutant of the RNA polymerase I

**DOI:** 10.1101/307199

**Authors:** Tommy Darrière, Michael Pilsl, Marie-Kerguelen Sarthou, Adrien Chauvier, Titouan Genty, Sylvain Audibert, Christophe Dez, Isabelle Léger-Silvestre, Christophe Normand, Anthony K. Henras, Marta Kwapisz, Olga Calvo, Carlos Fernández-Tornero, Herbert Tschochner, Olivier Gadal

**Author notes:** Corresponding author: (OG).

## Abstract

Most transcriptional activity of exponentially growing cells is carried out by the RNA Polymerase I (Pol I), which produces a ribosomal RNA (rRNA) precursor. In budding yeast, Pol I is a multimeric enzyme with 14 subunits. Among them, Rpa49 forms with Rpa34 a Pol I-specific heterodimer (homologous to PAF53/CAST heterodimer in human Pol I), which might be responsible for the specific functions of the Pol I. Previous studies provided insight in the involvement of Rpa49 in initiation, elongation, docking and releasing of Rrn3, an essential Pol I transcription factor. Here, we took advantage of the spontaneous occurrence of extragenic suppressors of the growth defect of the *rpa49* null mutant to better understand the activity of Pol I. Combining genetic approaches, biochemical analysis of rRNA synthesis and investigation of the transcription rate at the individual gene scale, we characterized mutated residues of the Pol I as novel extragenic suppressors of the growth defect caused by the absence of Rpa49. When mapped on the Pol I structure, most of these mutations cluster within the jaw-lobe module, at an interface formed by the lobe in Rpa135 and the jaw made up of regions of Rpa190 and Rpa12. *In vivo*, the suppressor allele *RPA135-F301S* restores normal rRNA synthesis and increases Pol I density on rDNA genes when Rpa49 is absent. Growth of the Rpa135-F301S mutant is impaired when combined with exosome mutation *rrp6*Δ and it massively accumulates pre-rRNA. Moreover, Pol I bearing Rpa135-F301S is a hyper-active RNA polymerase in an *in vitro* tailed-template assay. We conclude that wild-type RNA polymerase I can be engineered to produce more rRNA *in vivo* and *in vitro*. We propose that the mutated area undergoes a conformational change that supports the DNA insertion into the cleft of the enzyme resulting in a super-active form of Pol I.

**Author summary:** The nuclear genome of eukaryotic cells is transcribed by three RNA polymerases. RNA polymerase I (Pol I) is a multimeric enzyme specialized in the synthesis of ribosomal RNA. Deregulation of the Pol I function is linked to the etiology of a broad range of human diseases. Understanding the Pol I activity and regulation represents therefore a major challenge. We chose the budding yeast *Saccharomyces cerevisiae* as a model, because Pol I transcription apparatus is genetically amenable in this organism. Analyses of phenotypic consequences of deletion/truncation of Pol I subunits-coding genes in yeast indeed provided insights into the activity and regulation of the enzyme. Here, we characterized mutations in Pol I that can alleviate the growth defect caused by the absence of Rpa49, one of the subunits composing this multi-protein enzyme. We mapped these mutations on the Pol I structure and found that they all cluster in a well-described structural element, the jaw-lobe module. Combining genetic and biochemical approaches, we showed that Pol I bearing one of these mutations in the Rpa135 subunit is able to produce more ribosomal RNA *in vivo* and *in vitro*. We propose that this super-activity is explained by structural rearrangement of the Pol I jaw/lobe interface.

## Introduction

The nuclear genome of eukaryotic cells is transcribed by three RNA polymerases [1]. RNA polymerase II (Pol II) transcribes most of the genome and is responsible for all messenger RNA production. RNA polymerases III and I are specialized in the synthesis of a limited number of transcripts. RNA polymerase III (Pol III) produces small structured RNAs, including tRNAs and the 5S ribosomal RNA. RNA polymerase I (Pol I) produces a single transcript, the large polycistronic precursor (35S pre-rRNA in yeast; 47S in human), which constitutes the first step of ribosome biogenesis. Pre-rRNA is then processed by multiple successive steps into the mature rRNAs (25S, 18S, and 5.8S in yeast; 28S, 18S and 5.8S in human). Despite producing a single transcript, Pol I is by far the most active eukaryotic RNA polymerase, responsible for up to 60% of the total transcriptional activity in exponentially growing cells [2]. The strongly transcribed rRNA genes can be visualized using the DNA spread method developed by Miller *et al*, 1969, in which the 35S rRNA genes (rDNA) exhibit a “Christmas tree” configuration, with up to 120 polymerases per transcribed gene [3].

The full subunit composition and structural data are now available for the three nuclear RNA polymerases of the budding yeast *Saccharomyces cerevisiae* [4,5][6,7]. Pol I contains a core of shared or homologous subunits that are largely conserved in eukaryotes and archaea, as for the other two nuclear RNA polymerases [8]. The two largest subunits (Rpa190 and Rpa135) form the DNA-binding cleft that carries the catalytic center. Rpb5, Rpb6, Rpb8, Rpb10, and Rpb12 are shared with Pol II and Pol III, whereas Rpc40 and Rpc19 are only shared with Pol III. This nine-subunit core is associated with the stalk, a structure formed in Pol I by the heterodimeric complex Rpa43/Rpa14, which is involved in docking the essential Rrn3 initiation transcription factor to the enzyme [9–12]. The Pol I-Rrn3 complex interacts with promoter bound factors, the core factor (CF), forming the initially transcribing complex (ITC) [13–16]. Additionally, Pol I and Pol III contain subunits that are functionally and structurally related to Pol II-specific basal transcription factors, called the “Built-in Transcription Factors” [17–19]. Their presence in Pol I and Pol III results in a higher number of subunits, from 12 subunits in Pol II, to 14 and 17 for Pol I and Pol III respectively, and correlates with substantial transcript production from a few genes [8]. The heterodimer formed by Rpa34 and the N-terminal domain of Rpa49 (Rpa49Nt) in Pol I (equivalent to Rpc53 and Rpc37 in Pol III) is related to the basal transcription factor TFIIF, and stimulates endogenous transcript cleavage activity [18,20,21]. Rpc34 in Pol III and the Rpa49 C-terminal domain (Rpa49Ct) bear a tandem winged helix domain similar to TFIIE, also named A49tWH [18,20]. Rpa49Ct binds upstream DNA [15,22] and is involved in initiation and elongation [18,23]. Finally, Rpa12 in Pol I and Rpc11 in Pol III harbour a C-terminal domain involved in stimulating endogenous transcript cleavage activity, similar to that of TFIIS for Pol II [17,24].

Yeast genetic studies of Pol III and Pol I “Built-in Transcription Factors” have revealed striking differences, despite their clear similarities. Each Pol III subunit is essential, but none of the Pol I “Built-in Transcription Factors” is required for cell growth. Lack of Rpa34 or invalidation of Rpa49Nt, by removing the TFIIF-like heterodimer, has no effect on growth *in vivo* [18,25,26]. In contrast, full or C-terminal deletion of *RPA49* leads to a strong growth defect at all temperatures, which is more severe below 25°C [18,26,27]. Full deletion of *RPA12* leads to a strong growth defect at 25 and 30°C, and is lethal at higher temperatures [28]. Lack of the C-terminal extension of Rpa12 abolishes stimulation of intrinsic cleavage, without any detectable growth defect [17,24]. Finally, yeast strains carrying the triple deletion of *RPA49*, *RPA34* and *RPA12* are viable, but accumulate the growth defects associated with each of the single mutants [25].

Pol I is functional in the absence of Rpa49, but shows well-documented initiation and elongation defects, both *in vivo* and *in vitro* [23,26,27,29–31]. Restoration of active rRNA synthesis, in the absence of Rpa49, has been used to identify factors involved in initiation and elongation, such as Hmo1 and Spt5 [26,30,32]. Here, we made use of the spontaneous occurrence of extragenic suppressors of the growth defect of the *rpa49* null mutant [27] to better understand the activity of Pol I. We showed that the suppressing phenotype was caused by specific point mutations in the two largest Pol I subunits, Rpa190 and Rpa135. We identified a small area around Phe301 in subunit Rpa135, at an interface formed by the lobe in Rpa135 and the jaw made up of regions of Rpa190 and Rpa12, where most mutations cluster. Characterizing the Rpa135-F301S allele, we showed in an *in vitro* assay, that such Pol I mutant is more active than the wild-type enzyme. *In vivo*, overproduction of rRNA by Pol I bearing the Rpa135-F301S mutation was observed in backgrounds where the nuclear exosome activity is impaired by *RRP6* deletion.

## Results

### Isolation of extragenic suppressor mutants of the growth defect in absence of Rpa49

We characterized extragenic suppressors of the *RPA49* deletion to better understand how cell growth is achieved in the absence of Rpa49. *RPA49* full-deletion mutants show a strong growth defect at 30°C and are unable to grow at 25°C. However, spontaneous suppressors have been previously observed [27]. We quantified the frequency of occurrence of individual clones able to grow at 25°C. There was a low frequency of colony occurrence, comparable with the spontaneous mutation rate of a single control gene (*CAN1*; < 5.10^−6^). We isolated more suppressors after irradiating the cells with UV light. UV irradiation, resulting in a survival rate of about 50%, increased the frequency of suppressor mutations by approximately 10-fold. We identified clones that grew at 25°C after three days and selected individual colonies, called SGR for Suppressor of Growth Defect of *RPA49* deletion, with various growth rates. We ranked SGR from 1 to 186 based on their growth rates at 25°C; *SGR1* had a growth rate comparable to the wild-type (WT) condition (Fig. 1A). We crossed the 186 SGR clones with a strain of the opposite mating type bearing the deletion of *RPA49* to obtain diploid cells homozygous for the *RPA49* deletion and heterozygous for each suppressor. The restoration of growth of the diploids at 25°C showed that all suppressor phenotypes obtained were fully or partially dominant. We focused on the most efficient suppressor clones, SGR1 and SGR2, and performed tetrad analysis to follow segregation of the observed suppression phenotype. Each suppressor phenotype was linked to a single locus in the genome and SGR mutants had no strong growth defect (SGR1 in Fig. 1A). We used global genomic mapping of SGR1 and SGR2, derived from “genetic interaction mapping” (GIM) methods [33] (Materials and Methods; S1 Fig), and found a genomic linkage with genes encoding the two largest Pol I subunits: *RPA135* for SGR1 and *RPA190* for SGR2 (S1 Fig). Sequencing of the genomic DNA revealed that SGR1 bears a double mutation, whereas SGR2 bears a single one (*RPA135-I218T/R379K* and *RPA190-A1557V* alleles, respectively). Furthermore, we identified an additional mutant, SGR3, in *RPA135* (*RPA135-R305L)*. The heterogeneity of the growth induced by strong UV mutagenesis prevented suppressor cloning from the 183 other SGR clones.

**Figure 1.**
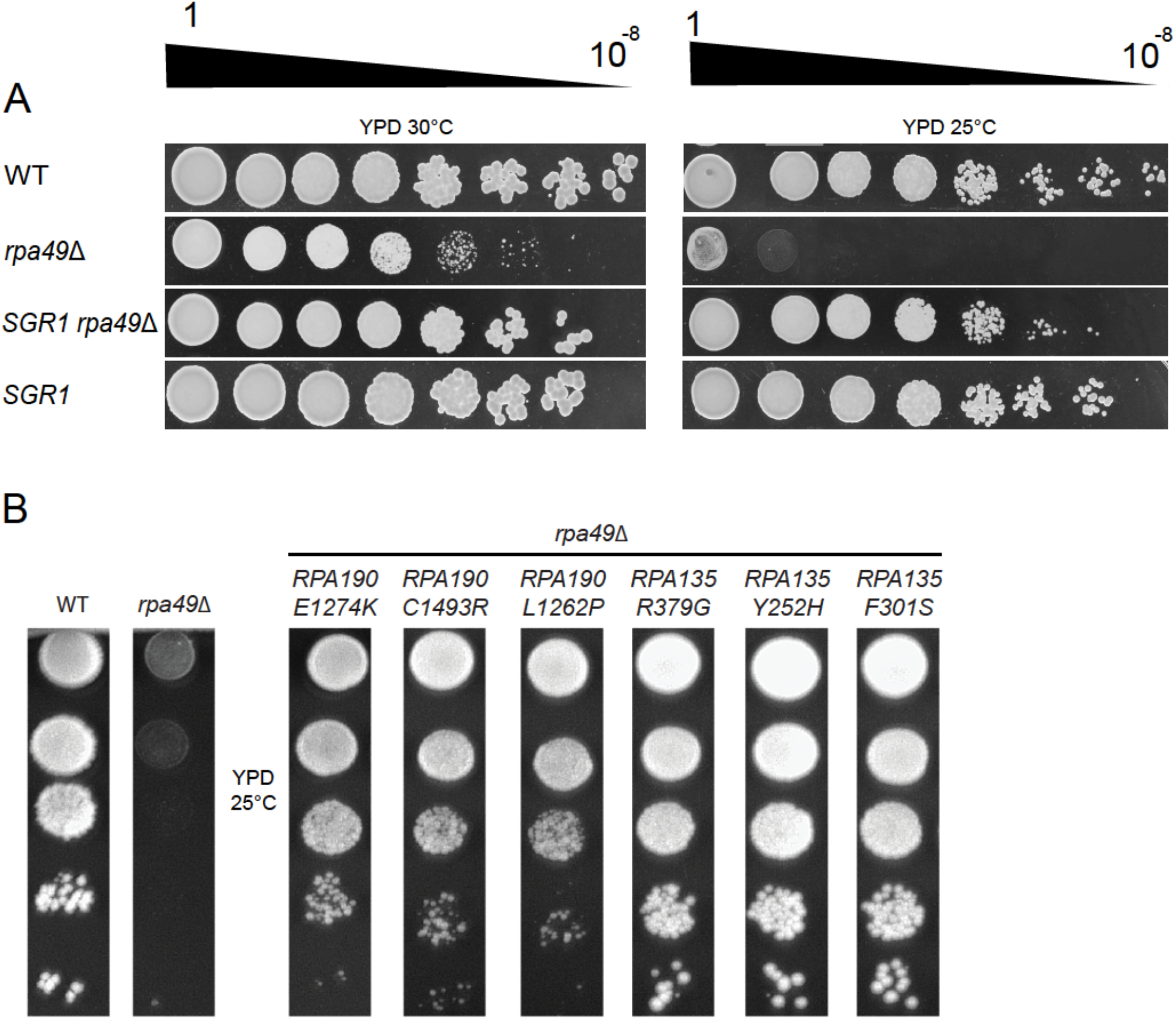
Alleles of *RPA190* and *RPA135* suppress the growth defect of the *rpa49*Δ mutant at various levels. (A) The *SGR1* mutant restores growth of the *rpa49*Δ mutant. Ten-fold serial dilutions of wild-type (WT), *rpa49*Δ single mutant, *SGR1* single mutant, and *SGR1/rpa49*Δ double mutant strains were spotted on rich media to assess growth at 30 and 25°C for three days. (B) Ten-fold dilutions of WT and *rpa49*Δ compared to *rpa49*Δ carrying various plasmids: pGL190_3 (*RPA190-E1274K*), pGL190_11 (*RPA190-C1493R*), pGL190_23 (*RPA190-L1262P*), pGL135_6prim (*RPA135-R379G*), pGL135_54 (*RPA135-Y252H*), or pGL135_33 (*RPA135-F301S*). Growth was evaluated after three days at 25°C. The strains and plasmids used are listed in S2-S3 Tables, respectively.

We next used the dominant phenotype of these suppressors to isolate more alleles, which suppress the deletion phenotype of *RPA49* in *RPA190* and *RPA135*. We constructed a library of randomly generated mutants (see Materials and Methods) by propagating plasmids bearing WT *RPA135* or *RPA190* in a mutagenic *E. coli* strain. After phenotypic selection of *rpa49*Δ mutants bearing a mutagenized Rpa190 or Rpa135 subunit at 25°C, each plasmid bearing a suppressor allele was extracted, sequenced, and re-transformed into yeast to confirm the suppressor phenotype. We thus isolated nine novel alleles of *RPA190* and thirteen of *RPA135* that were able to restore growth of *rpa49* deletion mutant at 25°C (S1 Table). We evaluated the suppression strength based on growth restoration relative to WT at 25°C, as for the SGR strains. Suppressor alleles obtained by mutagenesis of *RPA190* or *RPA135*, more effective than SGR1, 2, or 3 were identified (S1 Table). In conclusion, we identified 22 novel alleles of genes coding for the two largest Pol I subunits as extragenic suppressors of the *rpa49*Δ-associated growth defect.

### *rpa190* and *rpa135* mutant alleles can bypass the need of *RPA49* for optimal growth

The growth of the strains bearing one of six suppressor alleles (*RPA190-E1274K*, *RPA190-C1493R*, *RPA190-L1262P*, *RPA135-R379G*, *RPA135-Y252H*, and *RPA135-F301S*) was evaluated by a 10-fold dilution test (Fig. 1B), showing significant suppression by all in the absence of Rpa49. In previous genetic studies, other genetic backgrounds that alleviate the growth defect of *rpa49*Δ at 25°C were isolated: *rpa43-35,326* [26], decreased rDNA copy number [29], Hmo1 over-expression [30], or Spt5 truncations [32]. For all these mutants, rRNA synthesis was only partially restored in the absence of Rpa49 and significant transcription defects remained. Here, we focused on the *RPA135-F301S* allele, the most effective growth suppressor of the *RPA49* deletion: the *rpa49*Δ *RPA135-F301S* double mutant grew almost as well at 25°C as the WT strain (Fig. 1B).

We sought further insight into the effect of the suppressors by integrating the *RPA135-F301S* point mutation into the endogenous gene in three genetic backgrounds: WT, *rpa49*Δ (full deletion), or *rpa49*Δ*Ct*. Note that yeast bearing *rpa49*Δ*Ct* or *rpa49*Δ full-deletion have a similar growth defect, but have different Pol I subunits composition [26,27]. In the absence of Rpa49, Rpa34 does not associate with transcribing Pol I while in strains bearing the *rpa49*Δ*Ct* allele, Rpa34 and Rpa49Nt remain associated with the polymerase [26,27]. The growth rate was determined in each of these yeast strains at 30°C, in the presence or absence of *RPA135-F301S*. The suppressor allele *RPA135-F301S* had no effect on growth in the WT strain (doubling time of 102 min). The doubling time was 180 min for the *rpa49*Δ strain and *RPA135-F301S* restored growth to a doubling time of 135 min. We observed similar suppression on the *rpa49*Δ*Ct* background.

### Most suppressors mutations are clustered in the jaw-lobe module

Structural data are now available for Pol I in an inactive form [4,5], in complex with Rrn3 [12,23,34], associated with other initiation factors [13–15], in elongating forms [22,35], and in the paused state [36](Fig. 2A). We mapped Rpa135 and Rpa190 residues that suppress the growth defect of the mutant strain *rpa49*Δ onto the structure of WT Pol I in which the full structure of Rpa49 was determined [15] (Fig. 2B). Most of the suppressor mutations, which provided growth recovery (S1 table), appeared to be clustered at a specific interface between the two largest subunits, Rpa190 and Rpa135 (Fig. 2B), between the lobe (Rpa135 – salmon) and the jaw (Rpa190 – blue). In *RPA135*, we found five suppressor mutations, which modify a small region of 60 residues within the lobe domain (S2 Fig). Note that in this region, three amino acids “DSF” (D299, S300, F301), which are conserved among eukaryotic species, are all mutated in suppressors (S2 Fig). Substituted residues likely result in destabilization of this interface, suggesting a specific rearrangement of the interface lobe/jaw in each mutant. The jaw is also characterized by the presence of a *β*-strand in the structure of Rpa12 (Rpa12 – yellow: residues 46-51, Fig. 2B) that, along with four *β*-strands in Rpa190, forms a five-stranded anti-parallel *β*-sheet. The Rpa12 *β*-strand faces the Rpa135 lobe domain (residue 252 to 315 of Rpa135 – salmon), in which six independent mutations were found, including *RPA135-F301S*.

**Figure 2.**
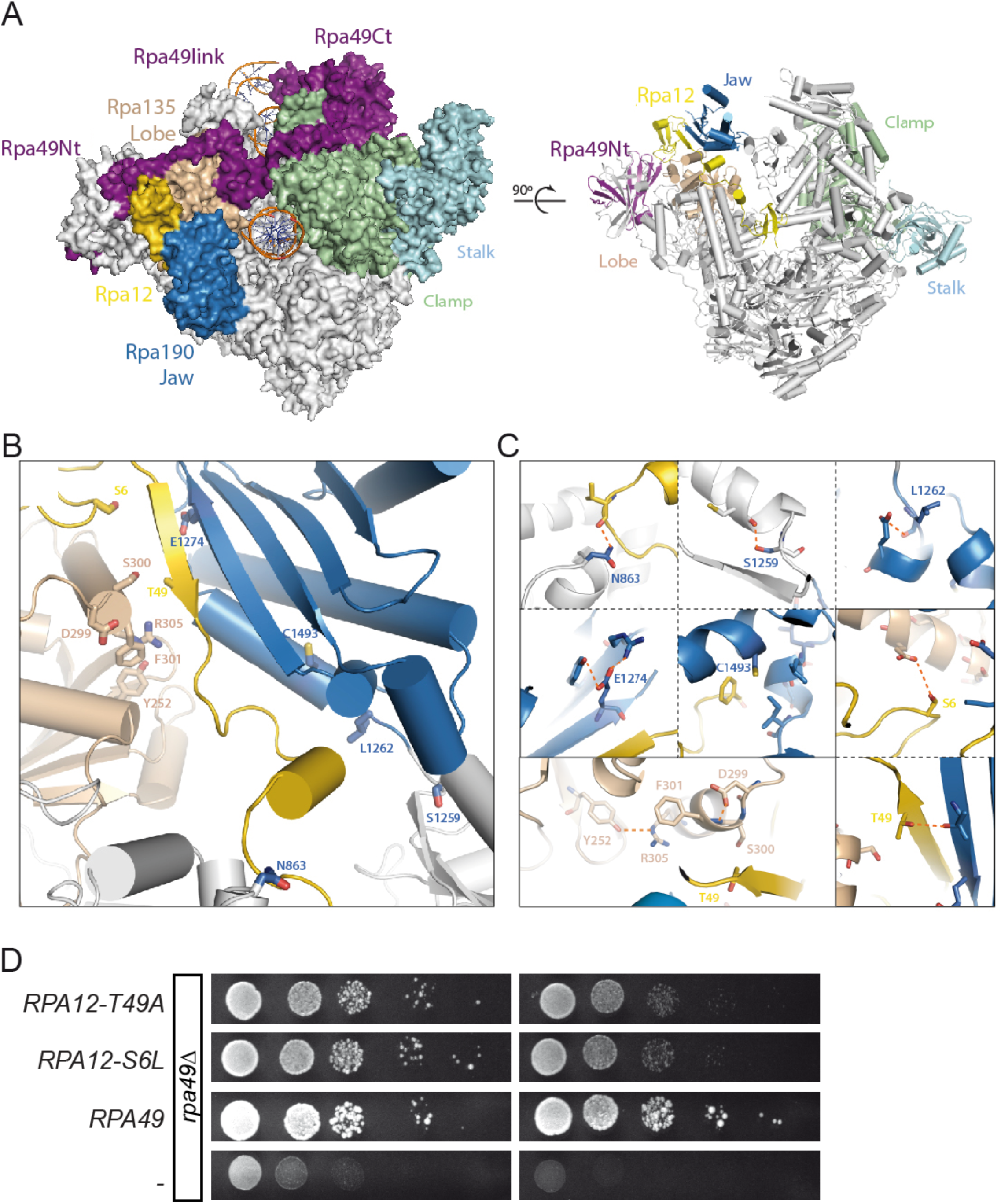
Mapping of the modified residues in Rpa190, Rpa135 on the structure of Pol I and isolation of Rpa12 alleles. (A) Two different views of the initially transcribing complex model and its 14 different subunits (PDB 5W66[15]). (B) Most mutated suppressor residues are clustered at the interface between the jaw (Rpa190, blue) and lobe (Rpa135, salmon) modules of Pol I. Note that residues 46-51 in Rpa12 (yellow) are part of this interface. (C) Zoom views of the areas containing the modified residues (Rpa190-N863, -S1259, -L1262, -E1274, -C1493; Rpa135-Y252, D299, S300, F301, R305; and Rpa12-S6, T49) shown in panel B. The figure was prepared with Pymol using the crystal structure of Pol I PDB 4C3I [4]). (D) Ten-fold dilutions of the *rpa49*Δ mutant carrying various plasmids: an empty pRS316 plasmid (-) YCp50-26 bearing *RPA49* (*RPA49*), pRS316-A12-S6L (*RPA12-S6L)*, or pRS316-A12-T49A (*RPA12-T49A)*. Growth was evaluated after three days at 25°C or two days at 30°C.

To evaluate the implication of Rpa12 in suppression, we then tested whether mutated alleles of *RPA12* could behave as suppressors. We generated a library of randomly mutagenized *RPA12*, and screen for *RPA12* alleles able to correct *rpa49*Δ growth defect at 25°C. Two dominant alleles (*RPA12-S6L* and *RPA12-T49A*) indeed efficiently suppressed the growth defect of *rpa49*Δ and of *rpa49*Δ*Ct* (Fig. 2D, not shown here for *rpa49*Δ*Ct*). *RPA12-S6L* and *RPA12-T49A* obtained by random mutagenesis are specifically located in the “hotspot” at the jaw/lobe interface. Threonine 49 of Rpa12 is located on the *β*-strand (Fig. 2C), facing residues D299, S300, and F301 of Rpa135, and Rpa190-E1274 (Fig. 2B). The second mutation, *RPA12-S6L*, is located in the N-terminal domain of Rpa12 (Fig. 2C).

In conclusion, all point mutations in Rpa190, Rpa135, and Rpa12 detected in the hotspot domain of the jaw/lobe interface substitute for *RPA49 in vivo*.

### Pol I bearing Rpa135-F301S or Rpa12-S6L restores efficient rRNA synthesis and Pol I occupancy on rRNA genes in the absence of Rpa49Ct *in vivo*

We used yeast mutant cells with a low (about 25 copies, +/− 3 copies) and stabilized (*fob1*Δ) number of rDNA repeats to better associate the growth phenotype of *RPA135-F301S* or *RPA12-S6L* allele to rRNA synthesis activity and Pol I density on transcribed genes *in vivo*. This genetic background is the best suited to study variations in the number of polymerase molecules per rRNA gene because it has a low number of rDNA copies, almost all in the active state with a very high Pol I loading rate [29,37,38]. We generated five strains in this low copy background (bearing single mutations; *rpa49*Δ*Ct*, *RPA135-F301S*, *RPA12-S6L* and double mutants combining *rpa49*Δ*Ct* with *RPA135-F301S* or *RPA12-S6L* alleles) and determined their doubling time (Fig. 3A) and *de novo* synthesis of rRNA (Fig. 3B). Note that rDNA copies number is similar between strains, as indicated by chromosome XII size in pulse-field gel electrophoresis (S3 Fig). The presence of the *RPA135-F301S* allele in Pol I effectively compensated the growth defect caused by the absence of C-terminal part of Rpa49 in this background.

**Figure 3.**
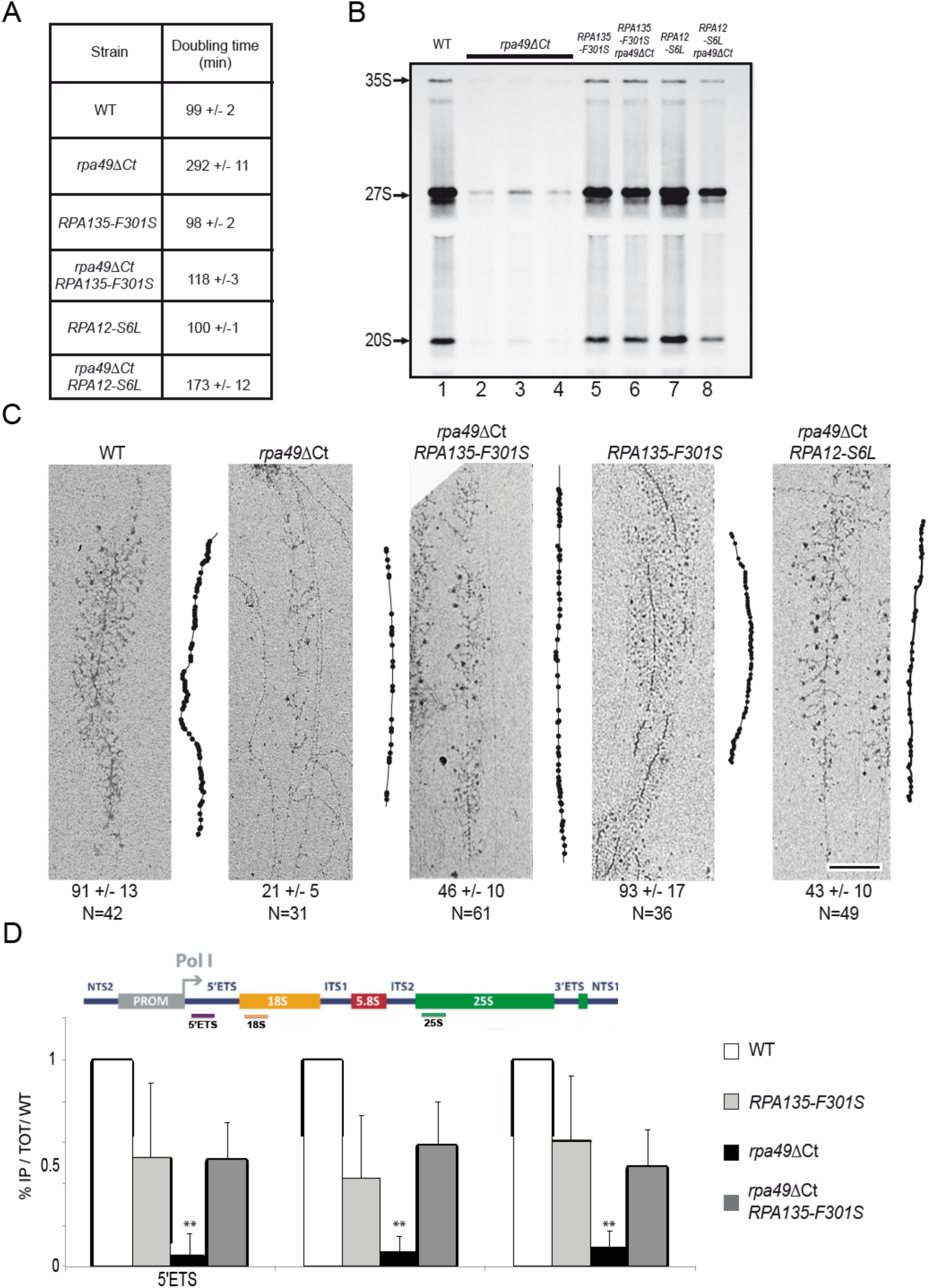
*RPA135-F301S* and *RPA12-S6L* alleles restore growth and rRNA synthesis, and modulate Pol I occupancy of rDNA genes in the absence of Rpa49Ct. (A) Doubling times of WT, *rpa49*Δ*Ct*, *RPA135-F301S*, *rpa49*Δ*Ct*/*RPA135-F301S* double mutant, *RPA12-S6L*, and the *rpa49*Δ*Ct*/*RPA12-S6L* double mutant in a low rDNA copy number background (see S2 Table). (B) *In vivo* labelling of newly synthesized RNAs. WT (lane 1), *rpa49*Δ*Ct* (lanes 2-4), *RPA135-F301S* (lane 5), the *rpa49*Δ*Ct*/*RPA135-F301S* double mutant (lane 6), *RPA12-S6L* (Lane 7), the *rpa49*Δ*Ct*/*RPA12-S6L* double mutant (lane 8) were grown to an OD_600_ of 0.8. Cells were then pulse-labeled with [8-^3^H] adenine for 2 min. Samples were collected, and total RNA extracted and separated by gel electrophoresis. (C) Representative Miller spreads of WT, *rpa49*Δ*Ct*, *RPA135-F301S*, *rpa49*Δ*Ct*/*RPA135-F301S*, and *rpa49*Δ*Ct*/*RPA12-S6L* double mutant. Panels on the right of each micrograph show interpretive tracing of the genes. Polymerases that appear on the gene are shown on the tracing by black dots. The number of polymerases counted on the genes is indicated below. N represents the number of individual spread genes used for quantification (see Materials and Methods). Scale bar = 200 nm. (D) ChIP analysis of Pol I occupancy at rDNA. Strains bearing WT Pol I, expressing Rpa49 lacking its C-terminal part (*rpa49*Δ*Ct*), bearing suppressor mutation (*RPA135-F301S*) or double mutant, were subjected to ChIP experiments using TAP tagged Rpa135, as described in Materials and Methods. Experiments were reproduced three times, representative Pol I occupancy relative to WT level are shown. Position of qPCR amplicons are depicted on rDNA unit. ** Marks significantly different values (p-values > 0.01) in student’s test of *rpa49*Δ*Ct* with WT, *RPA135-F301S* and double *rpa49*Δ*Ct RPA135-F301S*.

Labelling of the nascent rRNA was performed using a 2-min pulse with ^3^H adenine. We performed the labelling in three independent cultures because of heterogeneity due to random occurrence of suppressors in cell cultures of the *rpa49*Δ*Ct* mutant. When compared to a WT strain, RNA precursors synthesis was reduced approximately five-fold for *rpa49ΔCter*, even under permissive conditions (30°C) (compare Fig. 3B, lane 1 to lanes 2 to 4). Pol I activity in the presence of *RPA135-F301S*, with or without Rpa49Ct, was similar to that of the WT enzyme (Fig. 3B, lane 5 and 6). Similarly, *RPA12-S6L* partially restored Pol I activity in the absence of Rpa49Ct (compare Fig. 3B, lanes 7 and 8). Thus, *RPA135-F301S* and *RPA12-S6L* appeared to largely restore rRNA production in the absence of C-terminal part of Rpa49.

To get insight in Pol I activity in suppressors strains, we combined the rRNA synthesis quantification with the analysis of the Pol I distribution along the rRNA genes. We evaluated Pol I density on transcribed genes by performing Miller spreads, the only technique that currently allows the counting of individual Pol I molecules on a single rRNA genes [3,29]. Using Miller spreads, we previously showed that full deletion of *rpa49* resulted in a three-fold decrease of Pol I density per gene [3,29]. We show here that strain expressing the *rpa49*Δ*Ct* allele results in a four-fold decrease of Pol I density per gene, with about 21 Pol I detected per gene, as compared to about 91 detectable in WT condition (Fig. 2C). Expression of the *RPA135-F301S* or *RPA12-S6L* allele, WT for *RPA49*, had no detectable influence on Pol I density (Fig. 3C, *RPA135-F301S* and not shown). In contrast to strain *rpa49*Δ*Ct*, double mutant *rpa49*Δ*Ct RPA135-F301S* or *rpa49*Δ*Ct RPA12-S6L* showed significantly higher Pol I occupancy (46 and 43 respectively instead of 21 Pol I molecules per gene). Using ChIP, we reproduced that Pol I occupancy in absence of Rpa49Ct is drastically reduced [26]. We confirmed Miller spread quantification, in which *RPA135-F301S* significantly increased Pol I occupancy in absence of Rpa49Ct, although not to WT level (Fig. 3D).

Overall, these results show that the presence of the *RPA135-F301S*, or to a lesser extend *RPA12-S6L* allele, in a strain lacking C-terminal part of Rpa49 restores rRNA synthesis to WT levels. However, Pol I density on rRNA genes is only partly restored, indicative of an improved transcription initiation, or increased stability of elongating Pol I in absence of Rpa49Ct.

### Genetic interplay between suppressors alleles and Pol I domains

Extensive genetic characterization of Pol I subunits together with recent structural analysis have provided insight in their involvement in catalytic steps (initiation, pause release or termination). To investigate suppression mechanism, we then decided to explore which domains or subunits of Pol I are required for the suppression to occur. We tested deletion of *RPA14*, *RPA34* or *RPA12*, and of *RPA190* alleles (*rpa190*Δ*loop*) coding for Rpa190 lacking specific domain. The structure of Rpa190 revealed the presence of an extended loop inside the DNA-binding cleft folded in a “expander/DNA mimicking loop” conformation when Pol I is in an inactive, dimeric form [4,5]. This loop is inserted in the jaw domain of Rpa190 (Fig. 2B), in the vicinity of the mutation hotspot. A small deletion of this Rpa190 domain (1361-1390) resulted in a slight slow-growth phenotype [4]. We generated a larger deletion allele, *rpa190*Δ*loop* (deletion of residues 1342-1411 of Rpa190), and observed no associated growth defect (Fig. 4A). We were unable to generate a viable double mutant when combining this mutation with the *rpa49* full deletion. Thus, the DNA-mimicking loop is required for Pol I activity in the absence of Rpa49. We next tested whether deletion of this loop influences the suppression by the *RPA135-F301S* allele. Note that the *rpa190*Δ*loop* combined with *RPA135-F301S* had no growth phenotype. There was no difference in the growth of the *rpa49*Δ *RPA135-F301S* double mutant and that of the triple mutant *rpa49*Δ *RPA135-F301S rpa190*Δ*loop* (Fig. 4A). Thus, the expander/DNA mimicking loop of Rpa190 is not required for suppression, but is required for the viability of the *rpa49* deletion mutant.

**Figure 4.**
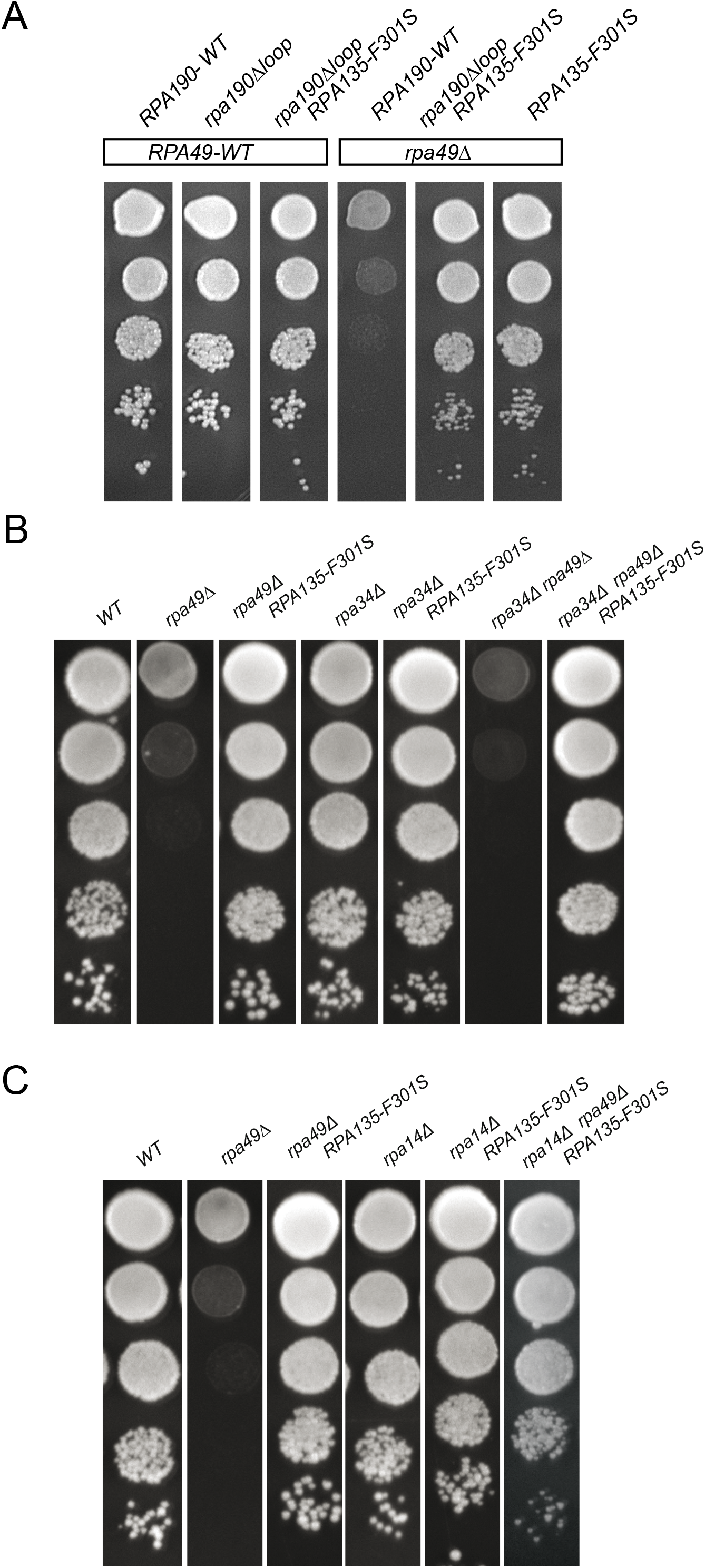
Rpa14, Rpa34, and the DNA mimicking loop of Rpa190 are not required for suppression. Deletion of the DNA mimicking loop of Rpa190 (A) or *RPA34* (B) does not modulate the suppression activity of *RPA135-F301S*. (C) *RPA135-F301S* suppresses the synthetic lethality between *rpa14*Δ and *rpa49*Δ. Ten-fold serial dilutions were performed and growth evaluated after three days at 25°C.

Rpa34 forms a heterodimer with Rpa49Nt, and Rpa14 is essential in absence of Rpa49. We then introduced *RPA135-F301S* in yeast strains lacking either Rpa34 or Rpa14 (Fig. 4B and C). Growth of *RPA135-F301S/rpa34*Δ and *RPA135-F301S/rpa14*Δ double mutants were not different from that of the single mutants. However, *RPA135-F301S* suppressed the growth defect of the viable double mutant, *rpa34*Δ *rpa49*Δ (Fig. 4B). The double deletion mutant lacking both Rpa49 and Rpa14 was not viable [25]. Introduction of the suppressor *RPA135-F301S*, by genetic crossing, resulted in a triple mutant (*rpa14*Δ *rpa49*Δ *RPA135-F301S*) that could grow, but slower than WT (Fig. 4C). We conclude that *RPA135-F301S* does not require Rpa14 or Rpa34 for the suppression to occur.

We next evaluated which part of Rpa12 subunit was required for the suppression to occur (Fig. 5). First, we evaluated growth of *rpa12* alleles when combined with *rpa49* deletion. The C-terminal region of Rpa12 (TFIIS-like) is inserted towards the active center of Pol I to stimulate intrinsic cleavage activity but is displaced during productive initiation and elongation steps. C-terminal deletion of Rpa12 resulted in normal growth [24] (Fig. 5A-lane 2), although the *rpa12*Δ*Ct* allele is unable to stimulate cleavage activity *in vitro* [17]. Full deletion of *RPA12* led to a heterogeneous growth phenotype when propagated at 30°C. To overcome this heterogeneity, we constructed a strain with *RPA12* under the control of the regulatable pGAL promoter. Depletion of Rpa12 on glucose containing medium, like full *RPA12* deletion, resulted in a slight growth defect at 25°C, which was stronger at 30°C [28] (Fig. 5A-lane 3). In contrast, *RPA49* deletion resulted in a growth defect at 30°C, which was stronger at 25°C [27] (Fig. 5A – lane 4). Combining *rpa12*Δ*Ct* with *rpa49*Δ resulted in a mild synergistic phenotype, with a stronger growth defect at both 25°C and 30°C (Fig. 5A – lane 5). The double mutant lacking both full Rpa12 and Rpa49 subunits was viable, but had a major growth defect [25] (Fig. 5A – lane 6).

**Figure 5.**
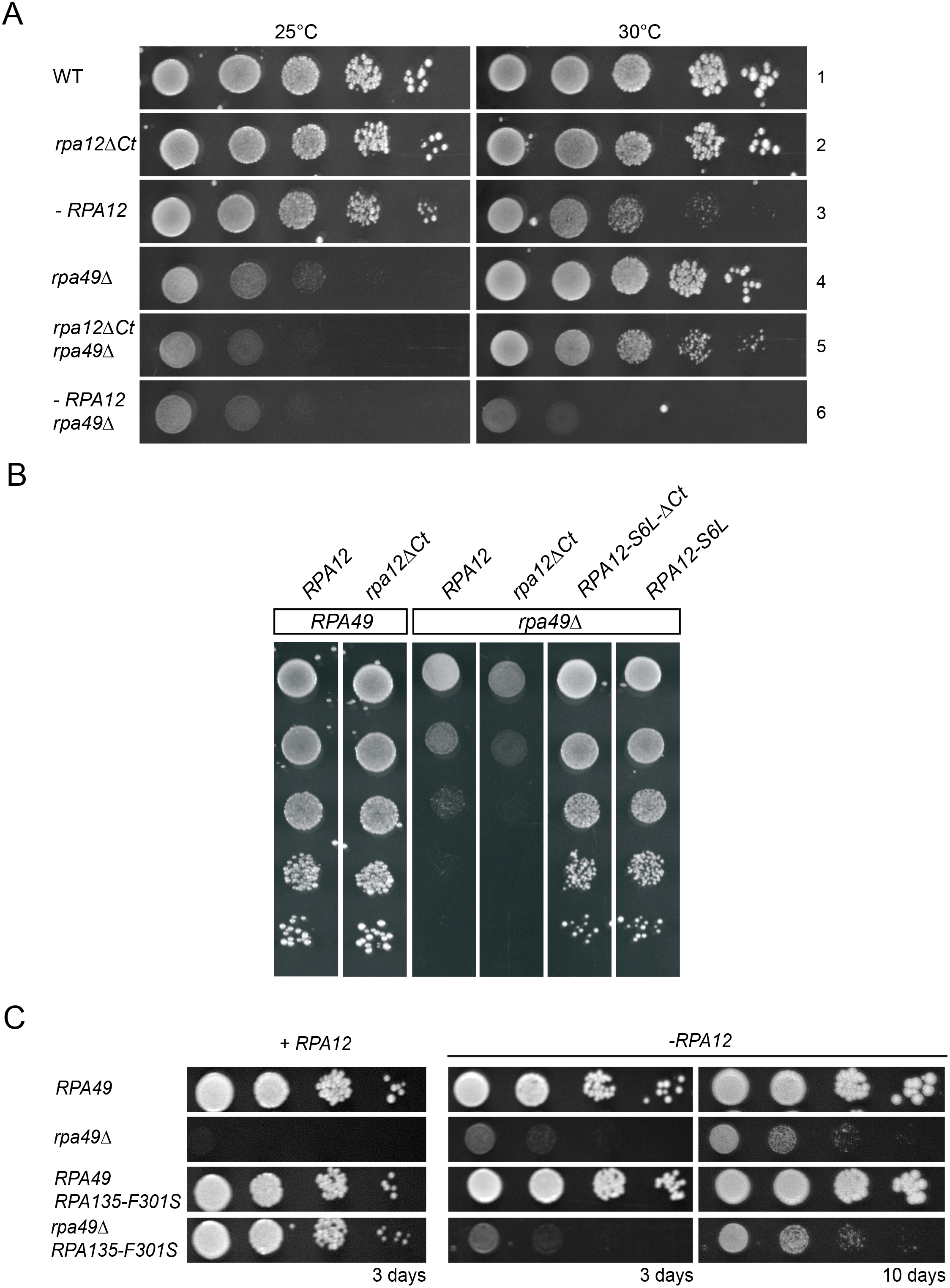
*RPA12* alleles can modulate the *rpa49*Δ-associated growth defect. (A) Growth of the double mutants: *rpa49*Δ *rpa12*Δ*Ct*, or *rpa49*Δ combined with full depletion of *rpa12*. Depletion of Rpa12 was achieved using a *pGAL-RPA12* construct on glucose containing medium (strain OGT30-1c). Ten-fold serial dilutions of OGT30-1c bearing pRS316-A12 (WT), pRS316-A12-DCt expressing Rpa12 bearing a C-terminal deletion of residues 65-125 (*rpa12*Δ*Ct*), or an empty plasmid pRS316 (*-rpa12*) were seeded onto media. The growth of *rpa49*Δ combined with *RPA12* depletion was tested using strain OGT30-3c bearing pRS316-A12 (*rpa49*Δ), pRS316-A12-DCt (*rpa12ΔCt rpa49*Δ), or an empty plasmid pRS316 (*-rpa12 rpa49*Δ). Growth was assessed after four days at 25°C or 30°C. (B) The C terminus of Rpa12 is not required for suppression. Ten-fold serial dilutions of OGT30-1c (*RPA49*-WT), bearing pRS316-A12 (WT) or pRS316-A12-DCter (*rpa12*Δ*Ct*), and OGT30-3c (*rpa49*Δ), bearing pRS316-A12 (WT), pRS316-A12-DCter (*rpa12*Δ*Ct*), pRS316-A12-S6L (*RPA12-S6L*), or pTD10 (*RPA12-S6L*-Δ*Ct*) were seeded onto media. Growth was assessed after four days at 25°C. (C) Suppression activity of *RPA135-F301S* is abolished in the absence of Rpa12. *RPA12*, under a regulatable promoter (pGAL) was either expressed (+*RPA12; left panel*) on galactose containing medium or repressed (−*RPA12; right panel*) on glucose containing medium. Ten-fold serial dilutions of *RPA49*, *rpa49*Δ, *RPA49*, *RPA135-F301S*, or *rpa49*Δ *RPA135-F301S* were grown at 25°C. Depletion of *RPA12* abolishes the suppression activity of *RPA135-F301S* (compare the left to the middle and right panels). Extended incubation (right panel, 10 days) was used to detect growth of the double mutant −*RPA12 rpa49*Δ on plates.

Secondly, in double *rpa12/rpa49* mutants, we tested the expression of the suppressors alleles, which were isolated (*RPA12-S6L*, *RPA12-T49A* or *RPA135-F301S*). We explored whether the C-terminal extension of Rpa12 was necessary for suppression of the *rpa49*Δ phenotype. We introduced the Rpa12 C-terminal truncation into the strain bearing both *rpa49*Δ and suppressor alleles *RPA12-S6L* or *RPA12-T49A*. *RPA12-S6L* and *RPA12-S6L*-Δ*Ct* resulted in similar suppression of the *rpa49*Δ growth defect (Fig. 5B). *RPA12-T49A* or *RPA12-T49A*-Δ*Ct* also suppressed equivalently (data not shown). We then introduced the *RPA135-F301S* allele in strains lacking Rpa49 with or without the entire Rpa12 subunit and assessed the suppression phenotype at 25°C (Fig. 5C). The growth defect of *rpa49*Δ was completely suppressed by the *RPA135-F301S* allele when Rpa12 was expressed (Fig. 5C, left panel), whereas suppression mediated by *RPA135-F301S* was not detected in the absence of Rpa12 (Fig. 5C, middle panel). After 10 days, *rpa49*Δ *rpa12*Δ double mutant behaved exactly the same with or without the *RPA135-F301S* allele, demonstrating that suppressor allele has no effect in absence of Rpa12 (Fig. 5C, right panel).

In conclusion, we show that *RPA135-F301S* suppression of *rpa49*Δ-associated growth defect does not require Rpa190 DNA mimicking loop, Rpa34, or Rpa14. Rpa12 C-terminal portion involved in stimulating cleavage activity is also not required for suppression. In contrast, Rpa12 N-terminal domain is required for the suppression to occur.

### *In vitro* characterization of RNA polymerase I bearing *RPA135-F301S*

*In vitro*, C-terminal part of Rpa49 is essential in promoter-dependent transcription assay [23]. Our *in vivo* analysis suggests that in *RPA135-F301S* mutant background, C-terminal part of Rpa49 is not required for rRNA synthesis. Our hypothesis is that Rpa135-F301S partly compensates the requirement for C-terminal part of Rpa49 in initiation. We used the promoter-dependent *in vitro* transcription system and tailed-template system to assess this hypothesis (Fig. 6A and B). Results were strikingly different when using promoter-dependent or tailed-template systems. After depletion of Rpa49, purified Pol I lacks both Rpa34 and Rpa49 subunits and is the so-called Pol A* complex [23,25,31]. Pol I lacking subunits Rpa49/Rpa34 (Pol A*, Fig. 6A lane 2) was almost inactive in promoter-dependent assay when compared to wild-type Pol I (WT) or Pol I bearing Rpa135-F301S (Fig. 6A, lane 1 and 3). Note that addition of recombinant Rpa34/Rpa49, Rpa49Ct alone or Rpa34/Rpa49N-ter stimulated transcription by Pol A* [23]. Ruling out our hypothesis, Pol I bearing Rpa135-F301S did not restore promoter-dependent activity of RNA Pol I lacking Rpa49 (Fig. 6A lane 4). We next tested RNA synthesis in a tailed-template system. Pol I lacking Rpa34/Rpa49 was partly deficient in tailed template assay (Fig. 6B, lane 2) [23]. Interestingly, in this promoter-independent assay, we clearly observed that the RNA synthesis by the polymerase bearing Rpa135-F301S was increased compared to one with the WT polymerase (Fig. 6B compare lane 1 to 3). Moreover, Pol I bearing Rpa135-F301S fully restored tailed template production of RNA Pol I lacking Rpa49 (Fig. 6B, lane 4). We conclude that in the *in vitro* promoter-dependent transcription assay, *RPA135-F301S* suppressor does not correct initiation defect due to the absence of Rpa49. However, in presence of Rpa135-F301S, a more efficient polymerase is engineered, able to produce more ribosomal RNAs from DNA tailed template.

**Figure 6.**
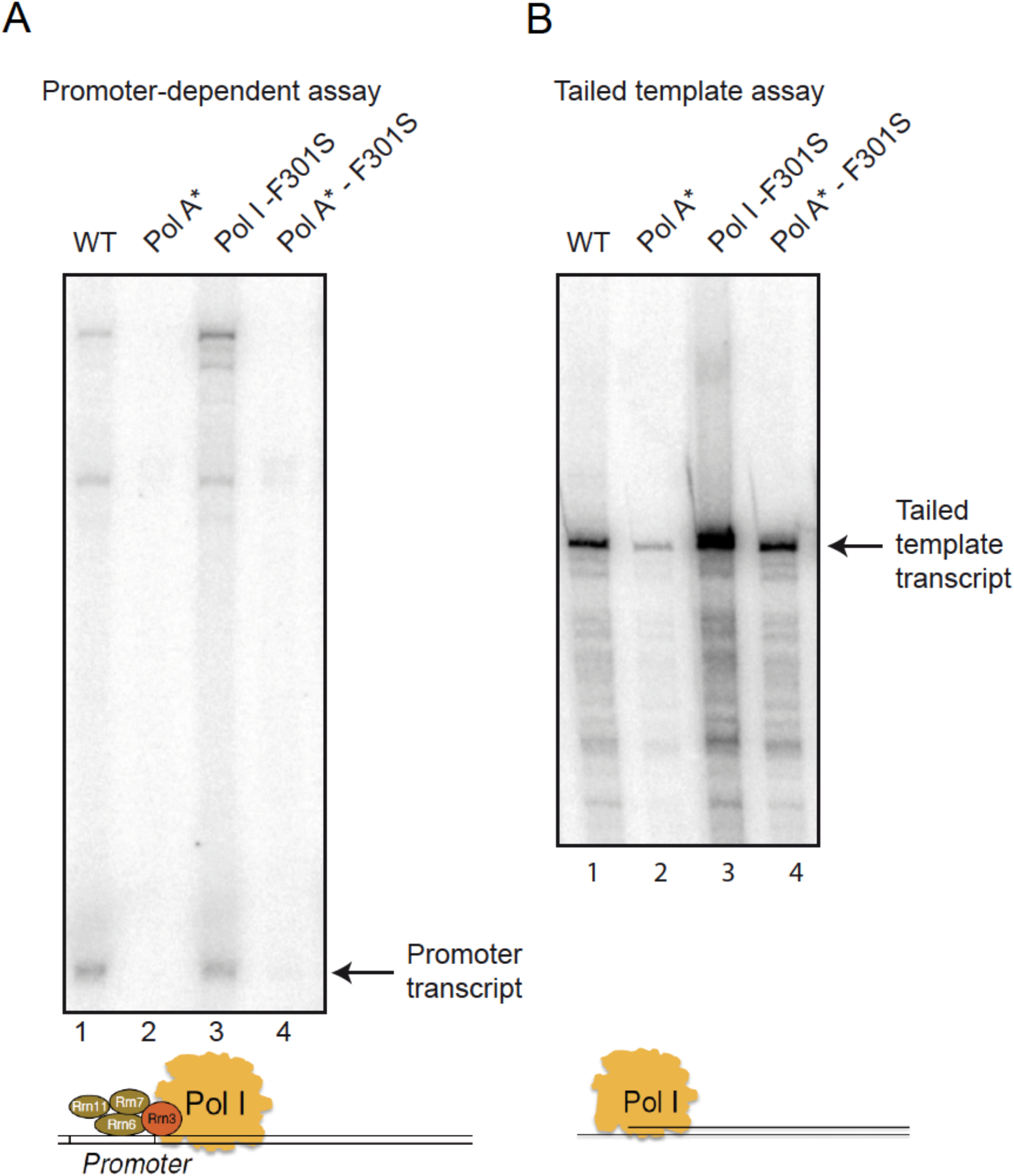
*In vitro* transcription assays of WT Pol I and Pol I mutants. WT, Pol A* (lacking Rpa34 and Rpa49), Pol I bearing Rpa135-F301S, and Pol A* bearing Rpa135-F301S were affinity-purified and 5 nM of each enzyme was used in either promoter dependent (A) or tailed template assay (B). Promoter-dependent assays were performed in the presence of 70 nM Rrn3 and 20 nM CF. Radiolabelled transcripts were separated on a denaturing polyacrylamide/urea gel and detected using a PhosphorImager. Note that upper radiolabeled bands in the experiment analyzing promoter-dependent transcription are due to nonspecific background labelling. Experiments were reproduced at least three times, with different Pol I concentration; representative experiments is shown.

### *In vivo* characterization of RNA polymerase I bearing Rpa135-F301S

*In vitro*, Pol I bearing Rpa135-F301S is over-producing RNA compared to WT. However *in vivo*, we could not reveal increased production of rRNA (2 min pulse labelling, see Fig. 3B). Pre-rRNAs which are not properly folded into pre-ribosome are targeted to degradation by the 3’ to 5’ exoribonucleolytic activity of the exosome [39]. We hypothesized that overproduced rRNAs in *RPA135-F301S* mutant background could be targeted by the nuclear exosome. Rrp6, part of the nuclear exosome complex, was deleted in a WT strain and in a strain bearing *RPA135-F301S* mutation. We observed a strong synergistic growth defect in strain bearing both *RPA135-F301S* and the deletion of *RRP6* (S5 Fig). Northern blot analysis (Fig. 7A) showed that accumulation in *RPA135-F301S* single mutant was indistinguishable from the WT for all RNA probed. *rrp6*Δ single mutant accumulates 23S and 35S (pre-)rRNA [40,41] and in correlation with the growth defect of the double mutant *RPA135-F301S rrp6*Δ, we could observe a 2-fold increase in 35S and 23S (pre-)rRNA accumulation as compared to *rrp6*Δ. Accumulation of 35S and 23S could indicate an increase of RNA production, or a defect in early maturation steps. We decided to directly assess over-expression of (pre-)rRNA using short *in vivo* labelling experiments (40 seconds). As previously reported with very short pulse labelling, accumulation of 20S rRNA is barely detectable, while 27SA and 35S are already accumulated [42]. We could detect a strong accumulation of newly synthesized (pre-) rRNA in the double mutant *RPA135-F301S rrp6*Δ compared to WT, *RPA135-F301S or rrp6*Δ strains. Note that increased background signal could indicate an accumulation of partially degraded, abortive transcripts or elongating rRNA transcript of various sizes. These results suggests that rRNA are over-expressed in strain bearing *RPA135-F301S*, but are quickly decayed by Rrp6. We confirmed this observation by evaluating ongoing transcription independently of decay machinery using high-resolution transcriptional run-on (TRO) analysis (Fig. 7C). Indeed, TRO assay make use of 10% sarkosyl, which permeabilizes cell membranes, reversibly blocks elongating polymerases and inhibits RNAse activity [43–46]. Permeabilized cells are then incubated with [*α*^32^P]-UTP to resume transcription. Neosynthesized radiolabeled RNAs are extracted, and used to probe slot-blots loaded with single strand DNA fragments complementary to rDNA locus. Using incorporation of [*α*^32^P]-UTP in the 5S rRNA transcribed by RNA polymerase III as internal control, TRO revealed a three-fold increase of rRNA transcription in cells bearing *RPA135-F301S* allele, irrespective of Rrp6 presence (Fig. 7C). These results confirmed that Pol I bearing Rpa135-F301S is over-producing RNA compared to WT and that over-produced RNAs are targeted for degradation by the exosome. All together, we concluded that Pol I bearing Rpa135-F301S is a hyper-active RNA polymerase *in vivo*.

**Figure 7.**
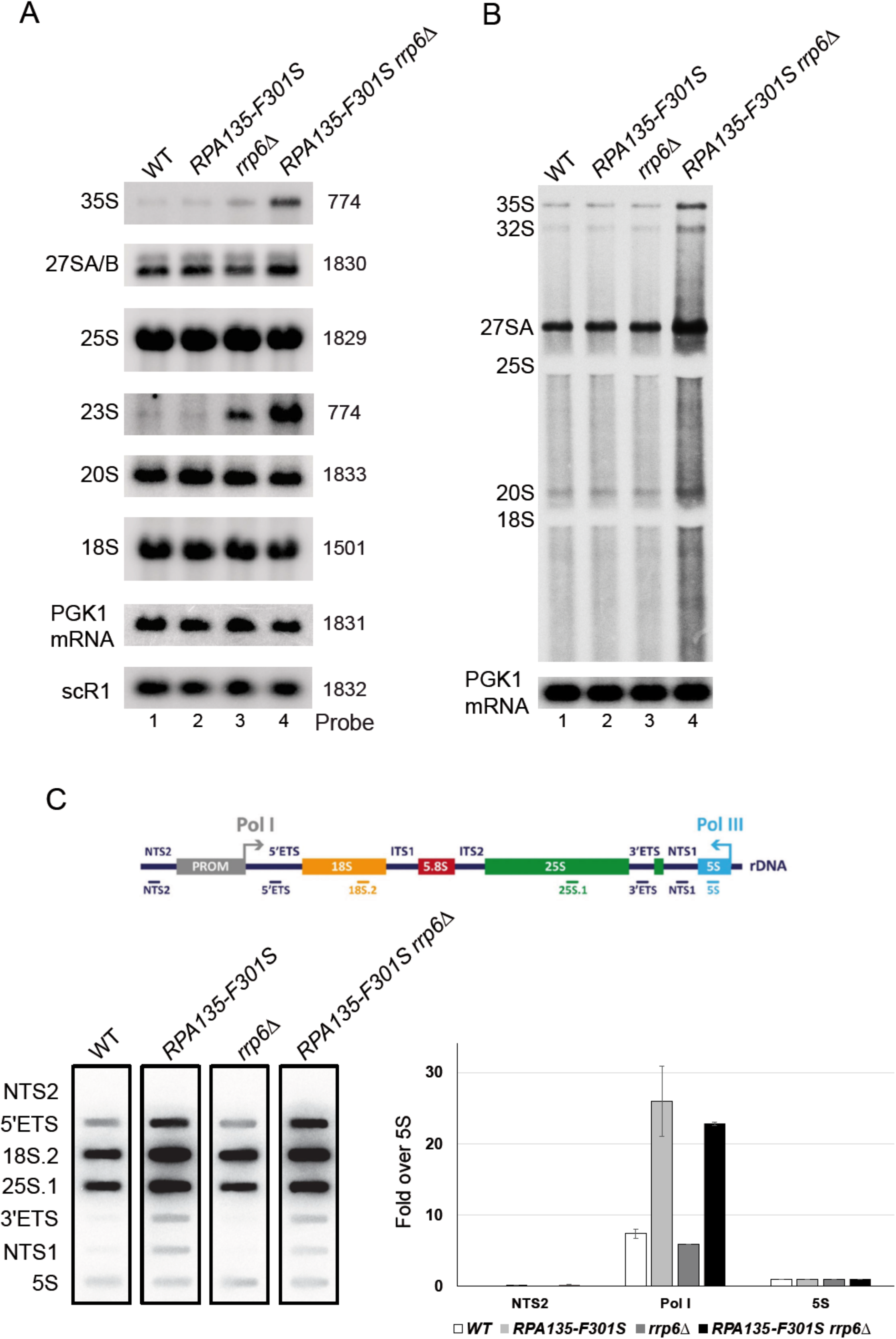
*RPA135-F301S* led to over-production of rRNA *in vivo*. (A) WT, *RPA135-F301S*, *rrp6*Δ and *RPA135-F301S*Δ *rrp6*Δ strains were grown to mid-log phase in glucose containing media. Cell samples were collected and total RNAs were extracted, separated by gel electrophoresis and transferred to a nylon membrane. The accumulation of the different (pre-) rRNAs was then analyzed by northern blot using different probes (see Materials and Methods). (B) *In vivo* labelling of newly-synthesized RNAs. WT, *RPA135-F301S*, *rrp6*Δ and *RPA135-F301S*Δ *rrp6*Δ strains were grown to an OD_600_ of 0.8. Cells were then pulse-labeled with [8-^3^H] adenine for 40 seconds. Samples were collected, and total RNA extracted and separated by gel electrophoresis. Newly-synthetized RNA are revealed by autoradiography, loading control was performed by northern blot (*PGK1* mRNA probe – 1831) on the same membrane. (C) High-resolution transcriptional run-on (TRO) analysis of WT, *RPA135-F301S*, *rrp6*Δ and *RPA135-F301S*Δ *rrp6*Δ strains. Nascent transcripts were labelled, and revealed using antisens oligonucleotides immobilized on slot-blot as described in Materials and Methods. Each experiment was performed twice; a representative example is shown in the lower left panel. NTS2, Pol I (mean of 5’ETS, 18S.2, 25S.1, 3’ ETS) are quantified relative to 5S signal in the lower right panel. Yeast rDNA unit is represented in the upper panel, with the position of the corresponding antisense oligonucleotides used.

## Discussion

Here, we characterized extragenic suppressors of the growth defect of the *rpa49* null mutant to better understand the activity of Pol I. We showed that altering a very specific area of Pol I resulted in an enzyme with modified catalytic properties sufficient to restore wild-type growth in absence of Rpa49.

### Suppressor mutations are not at the Rrn3-Pol I stalk interface

Our previous studies suggested the specific involvement of Rpa49 in the association and dissociation of initiation factor Rrn3 from the Pol I stalk [26,29]). Here, we show that genetically modified polymerases lacking Rpa49 or Rpa49Ct, with a single modified residue in Rpa190, Rpa135, or Rpa12, at a position diametrically to the position that binds to Rrn3, can initiate transcription and that strains harbouring them grow normally. Moreover, mutant Pol I with Rpa135-F301S does not restore promoter dependent activity in absence of Rpa49. We propose that, independently of the important interplay between Rpa49 and Rrn3 during initiation, Rpa135-F301S can stimulate Pol I activity.

### A novel role of Rpa12 subunit

The Rpa12 subunit is involved in stimulating the intrinsic cleavage activity of Pol I through a TFIIS-like domain at its C-terminus. Purified Pol I with Rpa12 lacking the C-terminal domain has no cleavage activity [17]. Furthermore, the C-terminal domain of Rpa12 can contact the active site of the polymerase in the inactive conformation and is retrieved in both initiation competent and elongating forms of the polymerase. However, the cleavage activity is re-activated when Pol I is paused [36]. Direct evidence that cleavage is not involved in suppression of the growth defect came from the experiments showing a fully functional suppressor phenotype for *RPA12*Δ*Ct-S6L*, which lacks the domain required for stimulating cleavage.

The N-terminal domain of Rpa12, at the surface of Pol I, is involved in the recruitment of the largest subunit, Rpa190 [24] and is required for docking this subunit to the enzyme. A linker region of Rpa12 connects its N-terminal module (equivalent to the N-terminal domain in the Pol II subunit Rpb9) at the surface of Pol I to its mobile C-terminal region (TFIIS-like) and is therefore indirectly required for cleavage. *In vitro*, purified Pol I that lacks Rpa12 has less activity than WT Pol I in promoter-dependent transcription assays (data not shown). Mutations in other Pol I domains, such as deletions in the Rpa190-DNA mimicking loop, Rpa34 or Rpa14, did not influence suppression of the *rpa49* deletion growth defect by the *RPA135-F301S* allele. In contrast, the Rpa12 linker was absolutely required for efficient suppression. Accordingly, *RPA135-F301S* allele was unable to restore efficient growth when Rpa12 was absent. Thus, Rpa12 and *RPA135-F301S* likely cooperate in the super-active Pol I enzyme. Note that rearrangement of Rpa12 have recently be shown to correlate with dissociation of Rpa49/Rpa34 heterodimer from Pol I, confirming the tight interplay between those subunits [47].

### Modification of the jaw/lobe interface may facilitate DNA cleft closure

Pol I undergoes major conformational changes during the transcription cycle, mainly affecting the width of the DNA-binding cleft [48]. During the initiation of transcription, the cleft aperture narrows from a semi-open configuration, as seen in cryo-EM structures of the enzyme bound to Rrn3 [12,23,34], to a fully closed conformation observed in transcribing complexes [13,15] (Fig. 8). This allows gripping of the transcription bubble inside the cleft (Fig. 8A-B). Following Rrn3 release, DNA binding is further secured, by the Rpa49-linker, which crosses the cleft from the lobe to the clamp, passing over the downstream DNA, and by the Rpa49Ct, which binds the upstream DNA in the vicinity of the clamp [15,22]. Therefore, Rpa49Ct is anchored in a position securing cleft closure (Fig. 8B).

**Figure 8.**
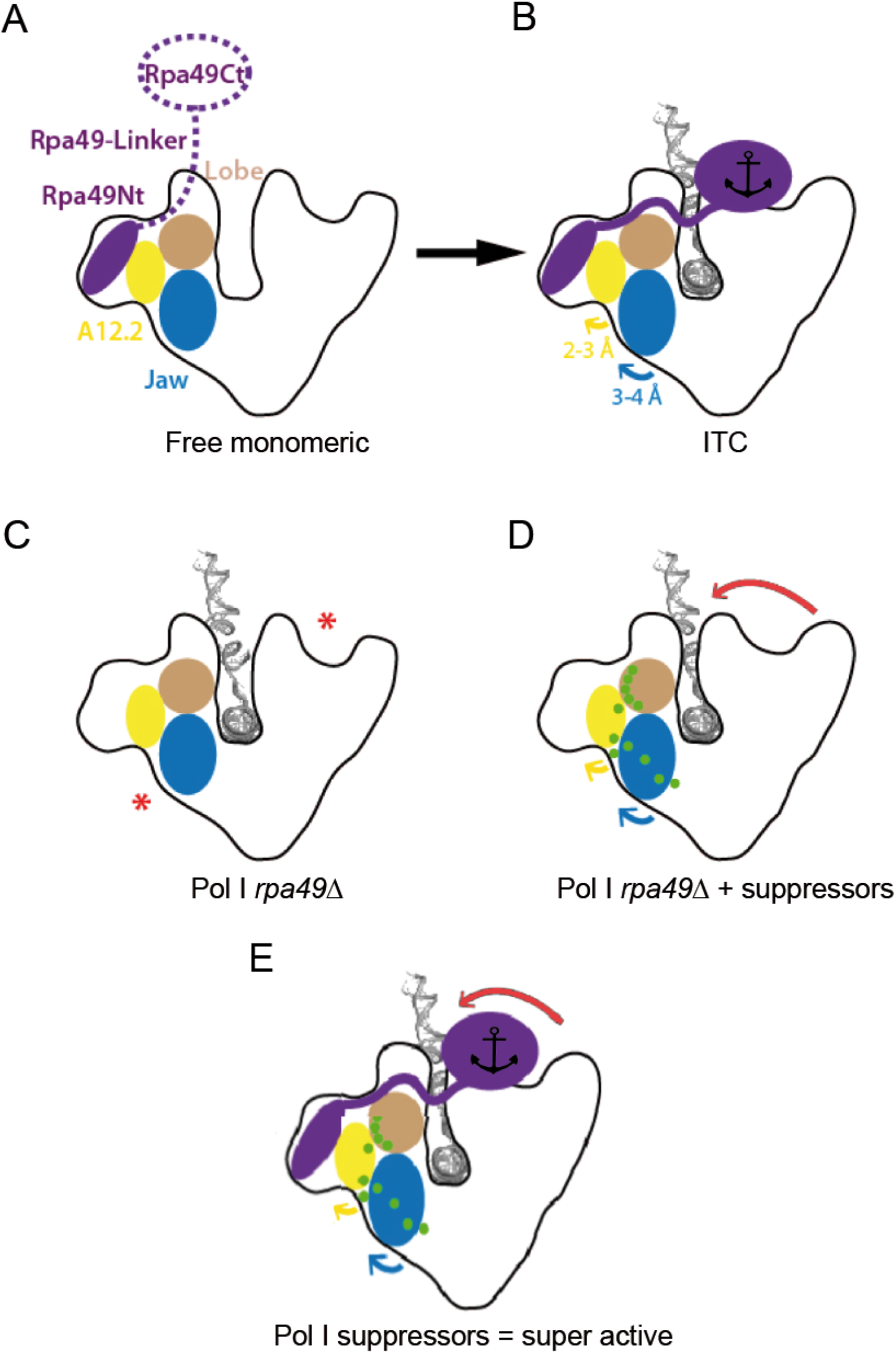
Schematic representation of Pol I. (A) Free monomeric Pol I with mobile Rpa49Ct and linker. (B) Initially transcribing complex (ITC) upon insertion of melted DNA in the presence of Rpa49 (purple). Rpa49Ct interacts with upstream DNA and the Rpa49-linker is folded, closing the cleft (black anchor). Movements of Rpa12 and the jaw with respect to the lobe are indicated with arrows. (C) Pol I lacking Rpa49 is likely defective in stabilizing the closed conformation in the DNA-binding cleft, resulting in a looser gripping of DNA inside the cleft (red asterisks). (D) Suppressor mutations (green) facilitate movement of the jaw/lobe interface and gripping of the DNA by the Pol I enzyme (red arrow), in the absence of Rpa49. (E) Combination of the presence of Rpa49 and a suppressor mutation (green dot) results in a super-active Pol I compared to the WT enzyme.

Cleft closure is achieved by the relative movement of two structural units, located on opposite sides of the cleft, pivot with respect to each other using five hinges [4]. The unit consisting of the shelf and clamp modules appears to be rigid, whereas the unit comprising the core and lobe modules, which is in the vicinity of the mutated residues involved in suppression of the *rpa49*Δ growth defect, undergoes internal rearrangements (S1 Movie). The most prominent reorganization within this latter unit affects the Rpa190 jaw domain, the outer rim of which shifts away from the DNA by approximately 3.7 Å, using the lobe/jaw interface as a hinge (Fig. 8B; blue arrow). This movement also involves the linker region of Rpa12, which contains a *β*-strand (residues 46-50) that completes a four-stranded *β*-sheet in the Rpa190 jaw domain. As a result, a short α-helix within the Rpa12 linker region shifts by approximately 3.0 Å (Fig. 8B; yellow arrow).

Rearrangements in the jaw made up of regions of Rpa190 and Rpa12 are likely essential to allow pivoting of the shelf-clamp unit against the core-lobe unit. Without such motion, cleft closure would be impossible (S1 Movie). Structural analysis suggests that the C-terminal domain of Rpa49 and its linker domain are involved in securing the closed cleft conformation [15,22]. Cleft closure is likely destabilized in the *rpa49*Δ mutant (Fig. 8C). We propose that Rpa135-F301S or Rpa12-S6L favors DNA capture by increasing the flexibility of the lobe/jaw/Rpa12 interface of Pol I relative to that of the WT polymerase, facilitating cleft closure in the absence of Rpa49 (Fig. 8D, red arrow). As shown *in vitro*, mutant Pol I with Rpa135-F301S is super-active. *In vivo*, the increased accumulation of pre-rRNA, detectable in absence of the nuclear exosome (*rrp6*Δ), is in agreement with the *in vitro* super-activity. We propose that Pol I bearing Rpa135-F301S might facilitate cleft closure, in addition to secure cleft closure by Rpa49. Such mutant enzyme could capture DNA more efficiently than WT polymerase (Fig. 8E). Alternatively, the mutations could also favour a ‘closed cleft’ conformation in the presence of DNA with little impact on flexibility.

### Pol I cleft closure is a limiting step in catalytic cycle

Here, we show that Pol I can be engineered to generate more rRNA. A super-active mutant of Pol II was already characterized: point mutation in trigger loop (Rpb1-E1103G) leads to increase RNA polymerization rate, affecting pausing and transcriptional fidelity [49,50]. Despite a very similar active site organization, an analogous mutation in the Pol I trigger loop resulted in a very different outcome, as it reduces the elongation rate [51]. Such observations led to the conclusion that Pol I catalytic cycle is very different than the one of Pol II, with different rate limiting steps [51–53]. We show here that mutations away of the active center can lead to a super-active form of Pol I. This difference can stem from wide-open configuration of the DNA-binding cleft in Pol I compared to other polymerases. We propose that Pol I cleft opening and closure is a limiting step in the Pol I catalytic cycle.

## Materials and Methods

### Construction of plasmids and yeast strains

The oligonucleotides used in this study are listed in S4 Table. Plasmids and details of the cloning steps are described in S3 Table. Randomly mutagenized *RPA190* and *RPA135* libraries were obtained by transformation and amplification of pVV190 and pNOY80, respectively, into XL1-red strains according to the manufacturer’s guidelines (XL1-Red Competent Cells, from Agilent Technologies). Yeast strains are listed in S2Table, and were constructed by meiotic crossing and DNA transformation [54][55]. The yeast media and genetic techniques were described previously [56]. yCNOD226-1a was obtained from yCNOD223-2a by switching the *KAN-MX* to the *NAT-MX* marker under the control of the MF(ALPHA)2/YGL089C promoter (alphaNAT-MX4), which allowed selection of MAT*α* haploid cells [33]. OGT9-6a is an offspring of yCNOD226-1a crossed with BY4741. OGT8-11a is an offspring of BY4742 crossed with Y1196. Strains OGT9-6a and OGT8-11a were plated on rich media and UV irradiated (5W/m^2^ during 5 second), resulting in 50% survival. LH514D and LH11D are suppressor clones of the growth defect selected from UV-irradiated OGT9-6a grown at 25°C. AH29R is a suppressor clone of the growth defect selected from UV-irradiated OGT8-11a grown at 25°C. Genetic interaction mapping (GIM) analysis of the *RPA49* deletion mutant was performed as described previously [33]. Microarray data were normalized using MATLAB (MathWorks, Inc., Natick, MA) as previously described [29]. OGT15-7b is an offspring of LH514D with BY4741, followed by homologous recombination using PCR-amplified fragments generated with oligos 1716 and 1717 and pCR4-HIS3 as template. yTD16-1a was first transformed with plasmid pCJPF4-GAL49-1. Then, *RPA135* was tagged by homologous recombination using PCR-amplified fragments generated with oligos 835 and 836 and genomic DNA of strain *RPA135-TAP* or yTD6-6c, generating yTD27-1 and yTD28-1a, respectively. Strain yTD25-1a bears a C-terminal deletion of *RPA49* generated by homologous recombination using PCR-amplified fragments generated with oligos 208 and 1515 and pFA6-KAN-MX6 as template. yTD11-1a was derived from strain yTD25-1a after switching to HPH-MX by homologous recombination using pUC19-HPH cut by *Bam*HI. C-terminal deletion of *RPA49 in* yTD29-1a and yTD30-1a were generated by homologous recombination using PCR-amplified fragments generated with oligos 649 and 650 and yTD11-1 genomic DNA as template, transformed into yTD27-1 and yTD28-1a, respectively. Genomic allelic insertion of *RPA12-S6L* in yTD31-1a and yTD23-1a was performed by homologous recombination using PCR-amplified fragments generated with oligos 1556 and 1557 and pRS316-A12-S6L-KAN as template, transformed into yTD27-1 and yTD29-1a, respectively.

TGT135-3b was obtained after sporulation of y27138 transformed by pNOY80. TGT135-3b and OGT15-9d were mated to generate TGT12. yTD2-3b and yTD2-3d are offspring of TGT12 transformed with pGL135_33. yTD6-6c and yTD6-6b were generated by homologous recombination using pTD2_6c_135TAP cut with *Xho*I-*Nsi*I, transformed into yTD2-3b and yTD2-3d, respectively. yTD48-1a was generated by deletion of the Rpa190 DNA-mimicking loop using homologous recombination with PCR-amplified fragments generated using oligos 1189 and 1194 and genomic DNA of strain SCOC2260 as template, transformed into BY4741.

yTD51-2c, yTD51-8a, and yTD51-5a are offspring of yTD48-1a mated with yTD37-3a. yTD36-2b is an offspring of yCN224-1a mated with yTD40-1a. yTD37-3d and yTD37-7d are offspring of yCN224-1a mated with yTD41-1a. yTD38-3d is an offspring of yCN225-1a mated with yTD40-1a and yTD39-8a is an offspring of yCN225-1a mated with yTD41-1a.

Strain yTD53-1a was constructed by homologous recombination using a PCR-amplified fragment generated with oligos 1634 and 1635 and pFA6a-KanMX6-GAL::3HA as template. OGT30-1a and OGT30-3a are offspring of yTD53-1a mated with yTD6-6b. yTD40-1a and yTD41-1a were generated by homologous recombination using PCR-amplified fragments generated with oligos 700 and 1679 and pFA6a-HA-KlURA3 as template, transformed into OGT30-3a and OGT30-1a respectively, switching *RPA135-TAP*-tag to untagged *RPA135*. Strain yMKS8-1a was constructed by homologous recombination using a PCR-amplified fragment generated with oligos 1711 and 1713 and pFA6a-KanMX6-GAL::3HA as template, transformed into strain yCD2-2a. yMKS9-9d is offspring of yMKS8-1a mated with yTD6-6c.

### Mapping extragenic suppressors allele by genetic linkage

Extragenic suppressors allele SGR were mapped using GIM methods, in which a strain is crossed with the entire pool of haploid deletion strains [57]. Here, deletions are used to evaluate linkage to SGR locus. Due to the genetic suppression, Individual deletions genetically linked to SGR are counter-selected in *rpa49*Δ background. Genetic mapping of SGR locus is based on strong genetic linkage of counter-selected deletion alleles (see S1 Fig), evaluated using micro-array of deletion bar-code (ArrayExpress accession E-MTAB-7831) [58].

### *In vivo* labelling and RNA extraction and analysis

Metabolic labelling of pre-rRNA was performed as previously described [59] with the following modifications. Strains were pre-grown in synthetic glucose-containing medium lacking adenine at 30°C to an OD_600_ of 0.8 at. One-milliliter cultures were labeled with 50 µCi [8-^3^H] adenine (NET06300 PerkinElmer) for 2 min. Cells were collected by centrifugation and the pellets were frozen in liquid nitrogen. RNA was then extracted as previously described [60] and precipitated with ethanol. For high molecular weight RNA analysis, 20% of the RNA was glyoxal denatured and resolved on a 1.2% agarose gel. Low molecular weight RNAs were resolved on 8% polyacrylamide/8.3 M urea gels.

### Miller spreads experiments and analysis

Chromatin spreading was mainly performed as described previously with minor modifications [61]. Carbon-coated grids were rendered hydrophilic by glow discharge instead of ethanol treatment. Negatively stained chromatin was obtained by short incubation with heavy metal followed by quick drying of the sample. Images were obtained using a JEOL JEM-1400 HC electron microscope (40 to 120 kV) with an Orius camera (11Mpixels). The position of the RNA polymerase I molecules and the rDNA fiber were determined by visual inspection of micrographs using Image J (http://rsb.info.nih.gov/ij/). Digital images were processed by software programs Image J and Adobe Photoshop® (v. CS6).

### *In vitro* promoter-dependent and tailed template transcription assays

*In vitro* promoter-dependent transcription reactions were performed as previously described [23,62] with some modifications. Briefly, 1.5 ml reaction tubes (Sarstedt safety seal) were placed on ice. Template (0.5–1 µl; 50–100 ng DNA) was added, corresponding to a final concentration of 5–10 nM per transcription reaction (25-µl reaction volume). Core factor (1–2 µl; 0.5 to 1 pmol/µl; final concentration 20–40 nM) and 1–3 µl Pol I (final concentration 4–12 nM) were added to each tube. Then, 20 mM HEPES/KOH pH 7.8 was added to a final volume of 12.5 µl. Transcription was started by adding 12.5 µl 2X transcription buffer. The samples were incubated at 24°C for 30 min at 400 rpm in a thermomixer.

25 nM of tailed templates were used in a total volume of 25 µl. The transcription was performed as described in promoter-dependent transcription reactions: Tailed templates are PCR-amplified fragments generated with oligos 1834 and 1835 and “pUC19tail_g-_601_elongated” as PCR-template, treated with Nb.BsmI (NEB) to generate a nick. Nicked site allows non-specific initiation of Pol I without the addition of initiation factors.

Transcription was stopped by adding 200 µl Proteinase K buffer (0.5 mg/ml Proteinase K in 0.3 M NaCl, 10 mM Tris/HCl pH 7.5, 5 mM EDTA, and 0.6% SDS) to the supernatant. The samples were incubated at 30°C for 15 min at 400 rpm in a thermomixer. Ethanol (700 µl) p.a. was added and the tubes mixed. Nucleic acids were precipitated at −20°C overnight or for 30 min at −80°C. The samples were centrifuged for 10 min at 12,000g and the supernatant removed. The precipitate was washed with 0.15 ml 70% ethanol. After centrifugation, the supernatant was removed and the pellets dried at 95°C for 2 min. RNA in the pellet was dissolved in 12 µl 80% formamide, 0.1 M TRIS-Borate-EDTA (TBE), 0.02% bromophenol blue, and 0.02% xylene cyanol. Samples were heated for 2 min with vigorous shaking at 95°C and briefly centrifuged. After loading on a 6% polyacrylamide gel containing 7M urea and 1X TBE, RNAs were separated by applying 25 watts for 30–40 min. The gel was rinsed in water for 10 min and dried for 30 min at 80°C using a vacuum dryer. Radiolabelled transcripts were visualised using a PhosphoImager.

### RNA extractions and Northern Hybridizations

RNA extractions and Northern hybridizations were performed as previously described [60]. For high molecular weight RNA analysis, 3µg of total RNA were glyoxal denatured, resolved on a 1.2% agarose gel and transferred to a nylon membrane. The sequences of oligonucleotides used to detect the RNA species: 35S rRNA, 25S rRNA, 20S rRNA, 18S rRNA, 27S rRNA, *PGK1* mRNA and *SCR1* ncRNA are respectively 774, 1829, 1833, 892, 1830, 1831 and 1832 are reported in S4 Table.

#### Chromatin immunoprecipitation (ChIP)

ChIPs were performed essentially as described previously [63]. At least three independent cultures of each yeast strain were grown to exponential phase OD_600_ = 0.4-0.8 in YPD (yeast extract-peptone-glucose) at 30°C, and cross-linked for 10 min at RT by the addition of formaldehyde (F1635, SIGMA) to a final concentration of 1.2%. Adding glycine quenched the cross-linking reaction. Cells were broken in lysis buffer (50 mM HEPES [pH 7.5], 150 mM NaCl, 1 mM EDTA, 1% Triton X-100, 0.1% deoxycholate Na, 0.1% SDS, 1 mM AEBSF [Euromedex] and cOmplete EDTA-free [Roche]) with glass beads (diameter, 0.425 to 0.6 mm; SIGMA) in a Precellys 24 homogenizer (bertin technologies) for 3 min, 6000 rpm at 4°C. Extracted chromatin was sonicated to obtain 300-500 bp DNA fragments. 100 µg of sonicated chromatin (protein content measured using BCA Protein Assay kit, ref# 23225, Thermo Scientific) was immunoprecipitated for 4h at 21°C on Pan mouse Dynabeads (Invitrogen, 200 µl). Immunoprecipitated DNA was purified and quantified by real-time qPCR using iTaq universal SYBR**®**Green Supermix (Bio-Rad) and the ViiA7 AB Applied Biosystems (Life Technologies). Primer pairs used for amplification are listed in S4 table. Signals were analysed with Quant Studio Real Time PCR Software v1.1 and are expressed as percentage of input DNA. Bars on the graph show the median of values normalised to wild-type value. Error bars correspond to standard deviations of at least three independent cultures for each strain.

#### Pulse-field gel electrophoresis (PFGE)

PFGE is a technique that resolves chromosome-sized DNA molecules in an agarose gel. Over-night YPD cultures at 30°C were harvested in 0.1% azide Na and kept on ice. Cells were washed twice in cold 0.05 M EDTA [pH 8.0]. ∼2.10^8^ cells were resuspended in 1ml of freshly prepared NZ buffer (Citrate Phosphate buffer 33 mM, ETDA 0.05 M [pH 8.0], Sorbitol 1.2 M). Cells were pellets (8000 rpm, 1 min, 4°C) and resuspended in 500 µL of Zymolyase buffer (NZ buffer with 20T zymolyase) and 500 µL LMP 2% (SeaKem LE agarose, Lonza, ref# 50001). Homogenized pellets were poured into the plug molds (∼100 µl/plug) and incubated for 30 min in humid chamber at 37°C then 10 min at 4°C. Solidified plugs were transferred in 14 ml tubes with round bottom and incubate with 400 µl/plug of PK buffer (EDTA 0.125 M [pH 9.5], Sarkosyl 1%, PK 1 mg/ml (Roche, ref# 03115801001) and incubated 1-2 hours at 37°C with gentle agitation. PK buffer was changed and plugs were incubated O/N with gentle agitation. De-proteinated plugs were washed with 10 ml TE followed by washing with 10 ml TE + 1 mM AEBSF (Euromedex) for 2 hours. AEBSF was washed out twice with 10 ml of 0.05 M EDTA [pH 8.0] for 30 minutes. Plugs were stored at 4°C till usage in TE or 0.05 M EDTA [pH 8.0]. Plugs were loaded on 0.8% agarose gel in 1×TAE buffer (Certified Megabase agarose, Bio-Rad, ref# 161-3109) and run in 1×TAE buffer for 50h in CHEF-DRIII System with chiller unit (Bio-Rad) with following parameters: 3V/cm, initial S/time 250 sec, final S/time 900 sec, angle 120. Gel was stained with ethidium bromide (10 mg/ml in water). Gel was processed for Southern blotting: DNA was transferred on nylon membrane by passive transfer in 10×SSC buffer (Amersham Hybond XL, GE Healthcare). Membranes were UV-cross-linked and hybridized overnight with ^32^P-labeled single-stranded DNA probes at 42 °C. Blots were washed twice with 2×SSC and 1% SDS for 20 min and once with 0.5×SSC and 1% SDS for 20 min at 37 °C. rDNA locus (chromosome XII) was detected with 18S specific probe (CATGGCTTAATCTTTGAGAC). This protocol was kindly provided by B. Pardo (IGH, Montpellier, France), and is extensively described in [64].

#### Transcriptional run-on analysis

TRO was performed as previously described [39,43]. Slot blots were loaded with single-stranded 80-mers DNA oligonucleotides: 1855 (NTS2), 1857 (5’ETS), 1859 (18S.2), 1860 (25S.1), 1861 (3’ETS), 1862 (NTS1), 1863 (5S US) and 1864 (5S DS).

## Acknowledgements

We would like to thank the imaging platform of Toulouse TRI for their assistance. We thank Margaux Cescato, Lilian Guillot and Jorge Perez-Fernandez for strain and plasmid constructions. YCp50-26 carrying *RPA49* was kindly provided by P. Thuriaux. We also thank Melanie Panarotto for the Miller spreads analysis. This work benefited from exchanges with all unit members, including Anthony Henras, Frederic Beckouët and Lise Dauban. We thank Stephanie Balor and Vanessa Soldan from the METI platform for the TEM acquisitions.

## Supporting information captions

**S1 Fig: Genetic interaction mapping (GIM) to identify suppressor mutations in SGR1 and SGR2.** (A) Schematic representation of GIM interaction assay. Enrichment ratios between control and SGR are used as read-out to map genetic interactions. (B) Relative enrichment of each barcode (black cross) along chromosome (in red) is used to map genetic linkage. Green curve represents mean in a sliding windows of 20 barcodes. Local maximum in such curve was used to identify locus of interest: RPA49 as positive control (upper panel, chr 14); RPA135 in SGR1 (middle panel, chr 16); RPA190 in SGR2 (lower panel, chr 15).

**S2 Fig: Mapping of the mutated residues in Rpa135**. (A) Structural domains of Rpa135 are depicted [4,5], including the lobe domain in which mutations are clustered. (B) Sequences alignment of positions 251-310 of Rpa135 from *S. cerevisiae* (NP_015335.1), compared with *S. pombe* (NP_595819.2), *H. sapiens* (NP_061887.2), *M. musculus* (NP_033112.2), *D. rerio* (NP_956812.2), *D. melanogaster* (NP_476708.1) and *C. elegans* (NP_492476.1). Color code from light to dark blue indicates residue conservations. Mutated residues at position 252 (Y to H), 299 (D to G), 300 (S to F), 301 (F to S or L) and 305 (R to L) are depicted in red.

**S3 Fig: Analysis of the rDNA size by pulse field gel electrophoresis.** Chromosomes from indicated strains (same as in Fig. 3D, two clones) were separated on the agarose gel and analysed by Southern blot using rDNA-specific probe.

S4 Fig: Promoter dependent *in vitro* transcription assays of Pol A* (lacking Rpa34 and Rpa49) and Pol A* bearing Rpa135-F301S are complemented with recombinant A34.5/A49 heterodimer. Promoter-dependent assays were performed as in fig. 6, with recombinant A49/A34.5 protein, described in [23].

S5 Fig: Ten-fold serial dilutions of *WT, RPA135-F301S, rrp6*Δ and *RPA135-F301S – rrp6*Δ grown at 25°C for 3 days.

**S1 Table**: List of 24 individual suppressor mutations of the growth defect of *rpa49*Δ strain in the Rpa190, Rpa135, and Rpa12 subunits.

**S2 Table**: Yeast strains used in this study.

**S3 Table**: Plasmids used in this study.

**S4 Table**: Oligonucleotides used in this study.

**S1 Movie**: Conformational changes in Pol I upon initiation. The movie starts with the closed cleft conformation (PDB 5W66 [15]), in which melted DNA occupies the cleft and then changes to the intermediate cleft conformation observed in monomeric Pol I (PDB 5M3M [35]). Relevant structural regions have been variously colored and labeled, and residues mutated in this report are shown in red.

