## Supplementary material for "Genetic analyses led to the discovery of a super-active mutant of the RNA polymerase I": Supp. figure and tables

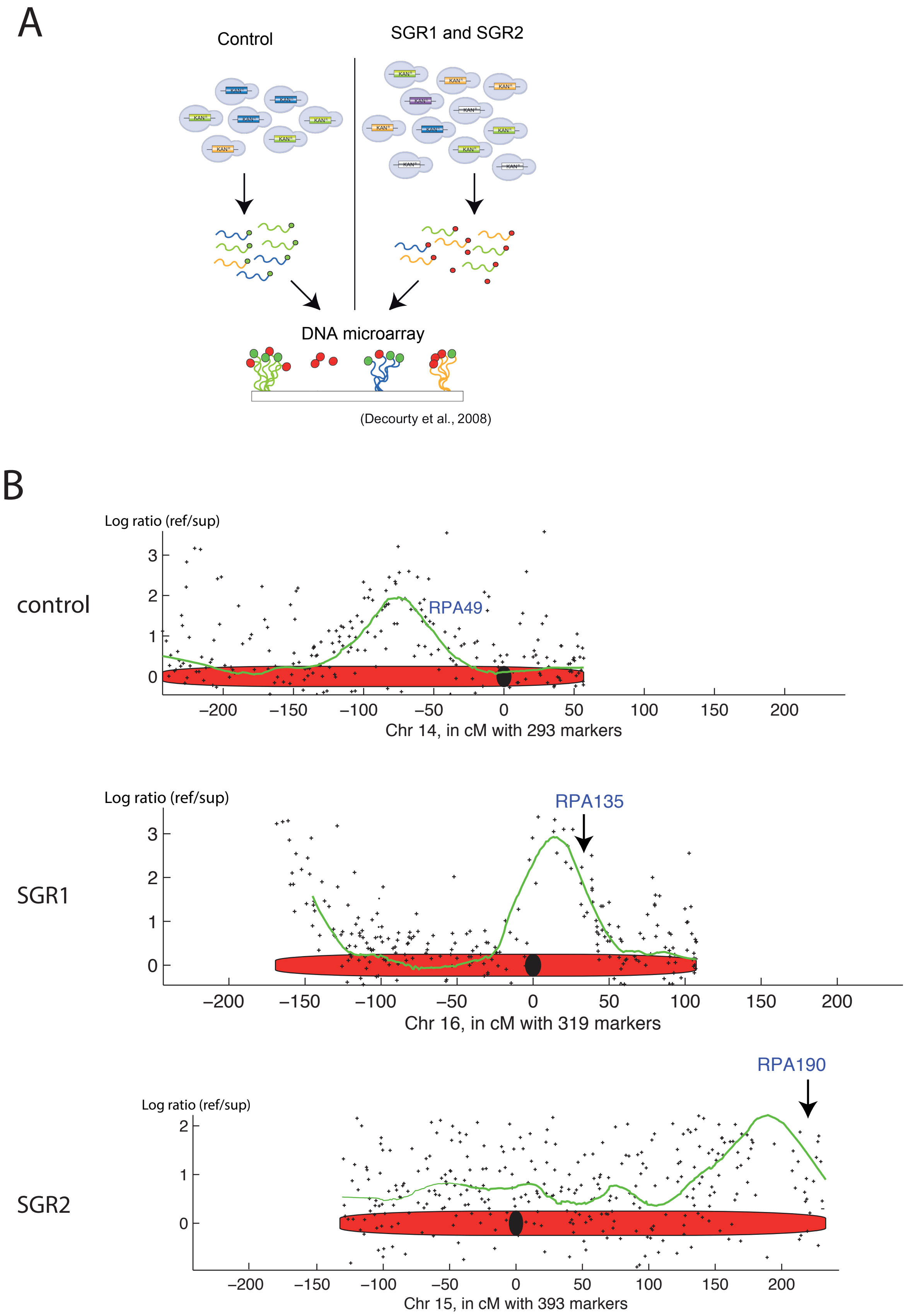


Genetic interaction mapping (GIM) to identify suppressor mutations in SGR1 and SGR2. (A) Schematic representation of GIM interaction assay. Enrichment ratio between control and SGR are used as read-out to map genetic interactions. (B) Relative enrichment of each barcode (black cross) along chromosome (in red) are used to map genetic linkage. Green curve represents mean in a sliding windows of 20 barcodes. Local maximum in such curve was used to identify locus of interest: RPA49 as positive control (upper panel, chr 14); RPA135 in SGR1 (middle panel, chr 16) ; RPA190 in SGR2 (lower panel, chr 15).

Supplementary Figure 1

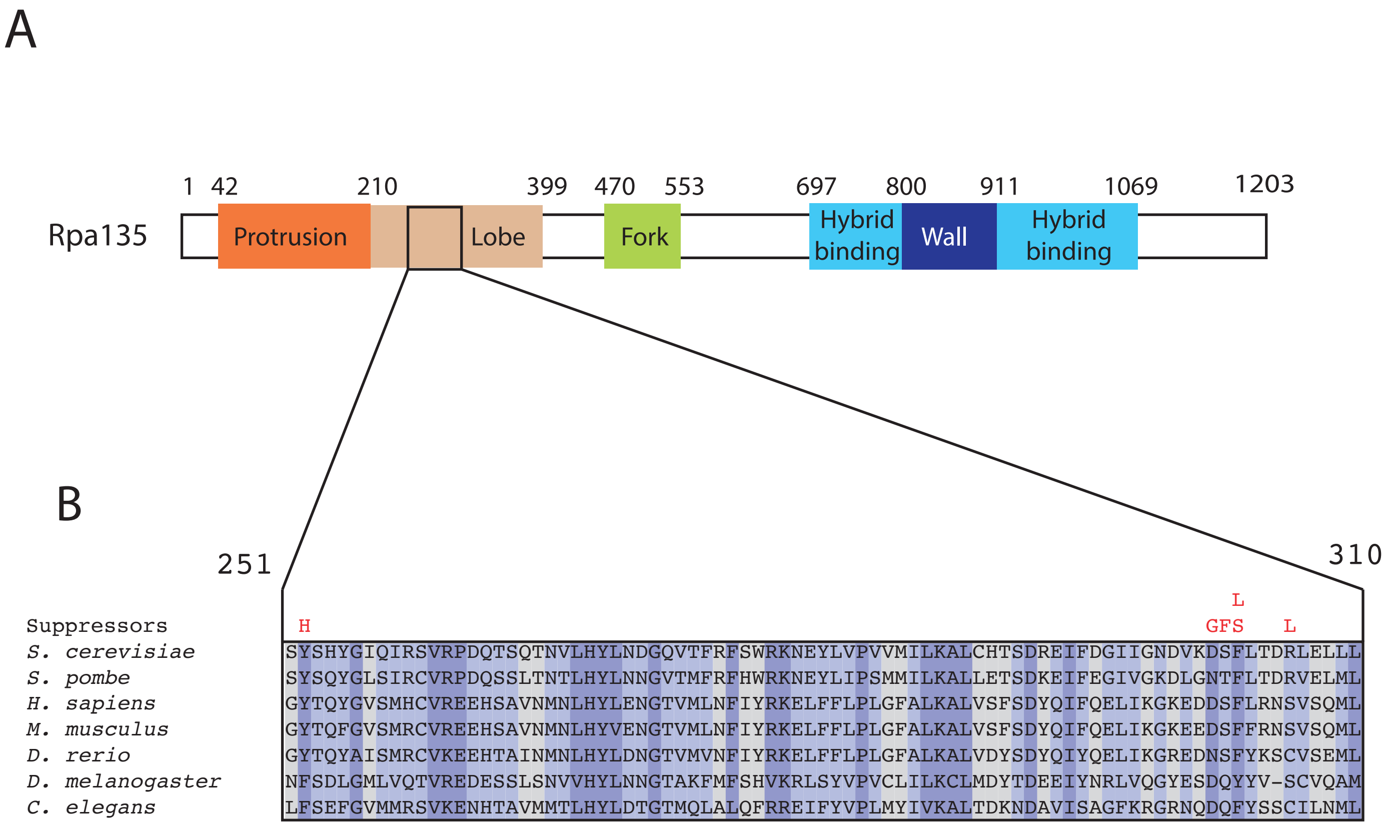


Mapping of the mutated residues in Rpa135. (A) Structural domains of Rpa135 are depicted [4,5], including the lobe domain in which mutations are clustered. (B) Sequences alignment of positions 251-310 of Rpa135 from *S. cerevisiae* (NP_015335.1), compared with *S. pombe* (NP_595819.2), *H. sapiens* (NP_061887.2) , *M. musculus* (NP_033112.2 ), *D. rerio* (NP_956812.2), *D. melanogaster* (NP_476708.1) and *C. elegans* (NP_492476.1 ). Color code from light to dark blue indicates residu conservations. Mutated residues at position 252 (Y to H), 299 (D to G), 300 (S to F), 301 (F to S or L) and 305 (R to L) are depicted in red.

Supplementary Figure 2

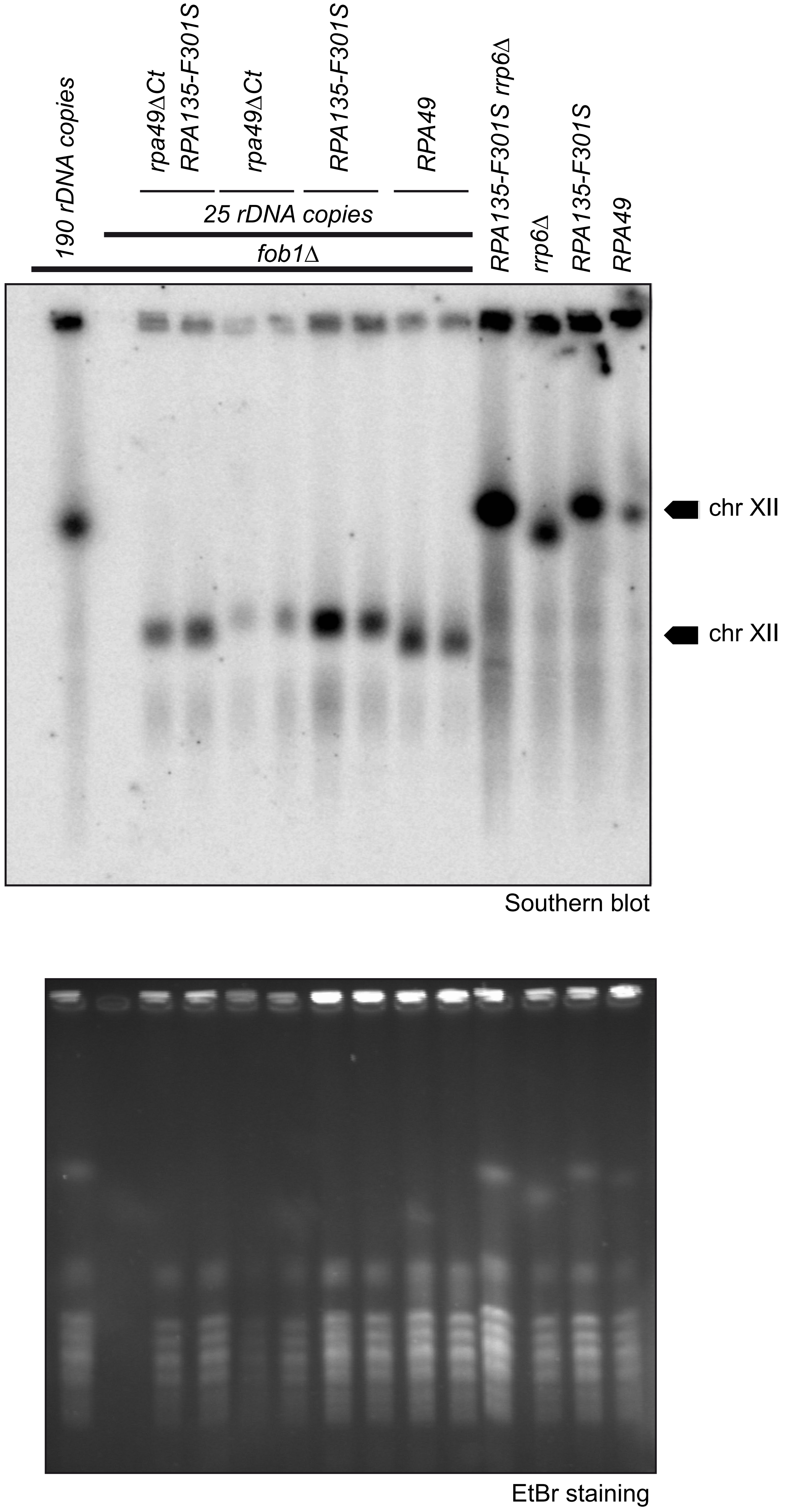


Analysis of the rDNA size by pulse field gel electrophoresis.
Chromosomes from indicated strains (same as in Fig. 3D, two clones) were separated on the agarose gel and analysed by Southern blot using rDNA-specific probe.

Supplementary Figure 3

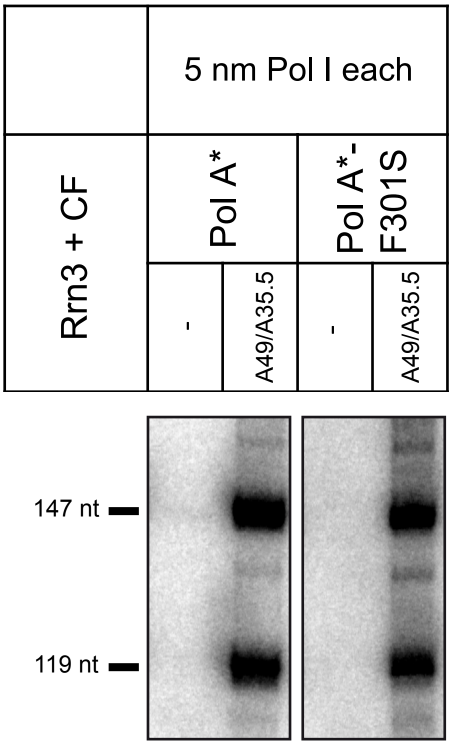


Promoter dependent in vitro transcription assays of Pol A* (lacking Rpa34 and Rpa49) and Pol A* bearing Rpa135-F301S are complemented with recombinant A34.5/A49 heterodimer. Promoter-dependent assays were performed as in fig. 6, with recombinant A49/A34.5 protein, described in [23].

Supplementary Figure 4

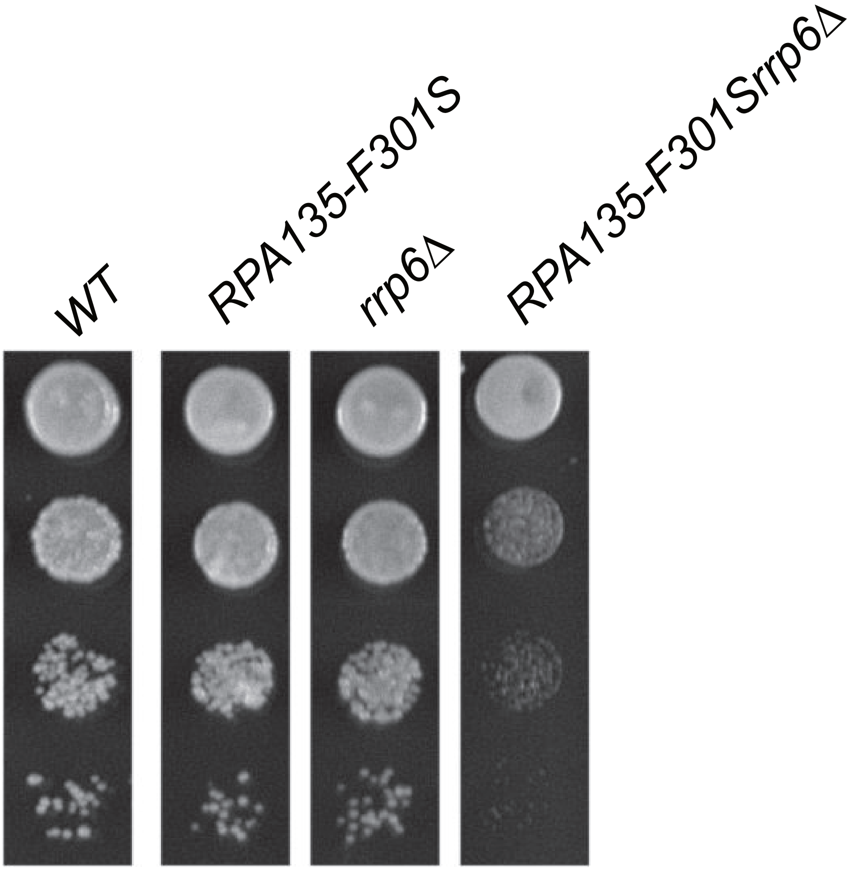


Ten-fold serial dilutions of WT, RPA135-F301S, rrp6Δ and RPA135-F301S rrp6Δ grown at 25°C for 3 days

Supplementary Figure 5

#### Supplementary tables

| Sub. | Allele | Strength | Hotspot | Location | Mutant effect |
|---|---|---|---|---|---|
| Rpa190 | N863T | Weak | Funnel | Funnel/ Rpa12-linker | Destabilization of Funnel/Rpa12 |
| Rpa190 | S1259L | Medium | Jaw | Jaw/Shelf hinge | Hinge conformation |
| Rpa190 | L1262P | Weak | Jaw | Jaw/Shelf hinge | Hinge conformation |
| Rpa190 | E1274K | Medium | Jaw | Jaw/ Rpa12-linker interface | Destabilization of Jaw/Rpa12 |
| Rpa190 | C1493R | Medium | Jaw | Jaw/ Rpa12-linker interface | Destabilization of Jaw/Rpa12 |
| Rpa135 | Y252H | Strong | Lobe | Jaw/Lobe interface | Destabilization of Jaw/Lobe |
| Rpa135 | D299G | Medium | Lobe | Jaw/Lobe interface | Destabilization of Jaw/Lobe |
| Rpa135 | S300F | Medium | Lobe | Jaw/Lobe interface | Destabilization of Jaw/Lobe |
| Rpa135 | F301S | Strong | Lobe | Jaw/Lobe interface | Destabilization of Jaw/Lobe |
| Rpa135 | F301L | Strong | Lobe | Jaw/Lobe interface | Destabilization of Jaw/Lobe |
| Rpa135 | **SGR3** R305L | Medium | Lobe | Jaw/Lobe interface | Destabilization of Jaw/Lobe |
| Rpa12 | S6L | Medium | N-terminal | Jaw/ Rpa12-linker interface | Destabilization of Jaw/Rpa12 |
| Rpa12 | T49A | Medium | Linker | Jaw/ Rpa12-linker interface | Destabilization of Jaw/Rpa12 |
| Rpa190 | L608S | Medium | | | |
| Rpa190 | E611K | Medium | | | |
| Rpa190 | S936A | Weak | | | |
| Rpa190 | **SGR2** A1557V | Weak | | | |
| Rpa135 | D157G | Medium | | | |
| Rpa135 | D157N | Medium | | | |
| Rpa135 | **SGR1** I218/ R379K | Medium | | | |
| Rpa135 | R379G | Strong | | | |
| Rpa135 | G580D | Medium | | | |
| Rpa135 | C584Y | Weak | | | |
| Rpa135 | I913V | Medium | | | |

**Supplementary table 1**. List of 24 individual suppressor mutations of the growth defect of *rpa49*Δ strain in the Rpa190, Rpa135, and Rpa12 subunits (Sub.). The suppressors were classified according to growth rate when combined with *rpa49*Δ mutant: weak, medium, or strong. SGR1, 2, and 3 depict alleles originally isolated after UV mutagenesis (see text). Thirteen of the 24 mutants, affecting 12 different positions, were found in a specific hot-spot shown in Figure 4. Residues substitution in the 3D structure were performed *in silico*, and putative mutant effects were deduced from the obtained structure.

#### Supplementary table 2: Yeast strains used in this study

| Referred to as | strain | Genotype | source |
|---|---|---|---|
| *WT in figures 1,4 and 7* | *BY4741* | *MATa his3Δ1 leu2Δ0 met15Δ0 ura3Δ0* | *Euroscarf* |
| | *BY4742* | *MATalpha his3Δ1 leu2Δ0 lys2Δ0 ura3Δ0* | *Euroscarf* |
| | *y1196* | *MATa his3Δ1 leu2Δ0 lys2Δ0 ura3Δ0 rpa49Δ::KANMX4* | *Euroscarf* |
| | *y27138* | *MATa/alpha his3Δ1/his3Δ1 leu2Δ0/leu2Δ0 ura3Δ0/ura3Δ0 lys2Δ0/LYS2 MET15/met15Δ0 rpa135Δ::KANMX4/RPA135* | *Euroscarf* |
| | *TGT135-3b* | *MATa his3Δ11 leu2Δ0 ura3Δ0 lys2Δ0 rpa135Δ::KANMX4 +pNOY80* | *Euroscarf* |
| | *TGT12* | *MATa/alpha his3Δ1/his3Δ1 leu2Δ0/leu2Δ0 ura3Δ0/ura3Δ0 lys2Δ0/lys2Δ0 rpa135Δ::KANMX4/RPA135 rpa49Δ::HPHMX4/RPA49* | *Euroscarf* |
| *rpa49Δ in figure 1, with the plasmids indicated in the legend* | *OGT9-6a* | *MATalpha his3Δ1 leu2Δ0 met15Δ0 ura3Δ0 rpa49Δ::alphaNATMX6* | *This study* |
| *rpa49Δ in figure 4A, with the plasmids indicated in the legend* | *OGT8-11a* | *MATa his3Δ1 leu2Δ0 lysΔ0 ura3Δ0 rpa49Δ::KANMX6* | *This study* |
| *SGR1 in figure 1* | *OGT15-7b* | *MATa his3Δ1 leu2Δ0 lysΔ0 ura3Δ0 HIS3::RPA135(I218T, R379K)* | *This study* |
| *SGR1 rpa49Δ in figure 1* | *LH514D* | *MATalpha his3Δ1 leu2Δ0 met15Δ0 ura3Δ0 rpa49Δ-alphaNAT RPA135(I218T, R379K)* | *This study* |
| | *AH29R* | *MATa his3Δ1 leu2Δ0 lysΔ0 ura3Δ rpa49Δ- kanmx4 RPA190-1557V* | *This study* |
| *SGR2* | *AC5-3b* | *MATalpha his3Δ1 leu2Δ0 lysΔ0 ura3Δ rpa49Δ-alphaNAT RPA190-1557V* | *This study* |
| *SGR3* | *LH11D* | *MATalpha his3Δ1 leu2Δ0 met15Δ0 ura3Δ0 rpa49Δ-alphaNAT RPA135-R305L* | *This study* |
| | *RPA135-TAP* | *MATalpha his3Δ1 leu2Δ0 met15Δ0 ura3Δ0 RPA135-TAP::HIS3MX6* | *202233825* [54] |
| | *yTD16-1a* | *MATa ade2-1 ura3-1 his3-11,15 trp1-1 leu2-3,112 can1-100 fob1Δ::NAT-MX, rDNA copy no. ~25* | *This study* |
| | *yTD6-6c* | *MATa ura3Δ0 his3-Δ1 leu2-Δ0 lys2-Δ0 RPA135-F301S-TAP-HIS3* | *This study* |
| *rpa49Δ in figures 2, 4A, 5* | *OGT15-9d* | *MATalpha his3Δ1 leu2Δ0 lysΔ0 ura3Δ0 rpa49Δ::HPHMX* | *This study* |
| *WT in figures 3* | *yTD27-1a* | *MATa ade2-1 ura3-1 his3-11,15 trp1-1 leu-3,112 can1-100 fob1Δ::NAT-MX RPA135-TAP-HIS3, rDNA copy number ~25 + pCJF4-LEU-GAL49* | *This study* |
| *RPA135-F301S in figure 3* | *yTD28-1a* | *MATa ade2-1 ura3-1 his3-11,15 trp1-1 leu-3,112 can1-100 fob1Δ::NAT-MX RPA135-F301S-TAP-HIS3, rDNA copy number ~25, + pCJF4-LEU-GAL49* | *This study* |
| | *yTD25-1a* | *MATa his3Δ1 leu2Δ0 met15Δ0 ura3Δ0 rpa49ΔC(186-416)::KAN-MX4* | *This study* |
| | *yTD11-1a* | *MATa his3Δ1 leu2Δ0 met15Δ0 ura3Δ0 rpa49ΔC(186-416)::HPH-MX4* | *This study* |
| *rpa49ΔCt in figure 3* | *yTD29-1a* | *MATa ade2-1 ura3-1 his3-11,15 trp1-1 leu-3,112 can1-100 fob1Δ::NAT-MX RPA135 -TAP-HIS3 rpa49ΔC(186-416)::HPH, rDNA copy number ~25, + pCJF4-LEU-GAL49* | *This study* |
| *rpa49ΔCt RPA135-F301S in figure 3* | *yTD30-1a* | *MATa ade2-1 ura3-1 his3-11,15 trp1-1 leu-3,112 can1-100 fob1Δ::NAT-MX RPA135-F301S-TAP-HIS3 rpa49ΔC(186-416)::HPH, rDNA copy number ~25, + pCJF4-LEU-GAL49* | *This study* |
| *RPA12-S6L in figures 3* | *yTD31-1a* | *MATa ade2-1 ura3-1 his3-11,15 trp1-1 leu-3,112 can1-100 fob1Δ::NAT-MX RPA135- TAP-HIS3 RPA12-S6L-KAN-MX, rDNA copy number ~25, + pCJF4-LEU-GAL49* | *This study* |
| *rpa49ΔCt RPA12-S6L in figure 3.* | *yTD23_1a* | *MATa ade2-1 ura3-1 his3-11,15 trp1-1 leu-3,112 can1-100 fob1Δ::NAT-MX RPA135- TAP-HIS3 RPA12-S6L-KAN-MX rpa49ΔC(186-416)::HPHMX, rDNA copy number ~25* | *This study* |

*Supplementary table 2 (continued)*

| | | | |
|---|---|---|---|
| | SCOC2260 | Mata ade2 arg4 leu2-3,112 trp1-289 ura3-52 RPB6::TAP-K.L.URA3 rpa190-ΔLOOP-KANMX | This study |
| rpa190Δloop in figure 4A | yTD48-1a | MATa his3Δ1 leu2Δ0 lysΔ0 ura3Δ0 rpa190Δloop::KANMX | This study |
| rpa190Δloop RPA135-F301S in figure 4A | yTD51-2c | MATalpha his3Δ1 leu2Δ0 met15Δ0 ura3Δ0 lys2Δ0 rpa190Δloop::KANMX RPA135-F301S-URA3Kl | This study |
| rpa190Δloop RPA135-F301S rpa49Δ in figure 4A | yTD51-8a | MATa his3Δ1 leu2Δ0 lys2Δ0 met15Δ0 ura3Δ0 RPA135-F301S-URA3Kl rpa49Δ::HPHMX | This study |
| RPA135-F301S rpa49Δ in figure 4A | yTD51-5a | MATa his3Δ1 leu2Δ0 lysΔ0 ura3Δ0 RPA135-F301S-URA3Kl rpa49Δ::HPHMX | This study |
| rpa49Δ in figures 4B and 4C | yCN223-2a | MATalpha his3Δ1 leu2Δ0 met15Δ0 ura3Δ0 lys2Δ0 rpa49Δ::KANMX | This study |
| | yTD2-3b | MATa ura3Δ0 his3-Δ1 leu2-Δ0 lys2-Δ0 rpa135::KANMX, pGL135_33 (ARS/CEN URA3 RPA135-F301S) | This study |
| rpa49Δ RPA135-F301S in figures 4B and 4C | yTD2-3d | MATa his3Δ1 leu2Δ ura3Δ0 lys2Δ0 rpa135Δ::KANMX rpa49Δ::HPHMX + pGL135_33 (ARS/CEN RPA135-F301S) | This study |
| rpa34Δ in figure 4B | yCN224-1a | MATalpha his3Δ1 leu2Δ0 met15Δ0 ura3Δ0 lys2Δ0 rpa34Δ::KANMX | This study |
| rpa34Δ RPA135-F301S in figure 4B | yTD36-2b | MATalpha his3Δ1 leu2Δ0 ura3Δ0 lys2Δ0 rpa34Δ::NATMX RPA135-F301S::URA3 | This study |
| rpa34Δ rpa49Δ in figure 4B | yTD37-7d | MATalpha his3Δ1 leu2Δ0 ura3Δ0 lys2Δ0 rpa34Δ::NATMX rpa49Δ::HPHMX | This study |
| rpa34Δ rpa49Δ RPA135-F301S in figure 4B | yTD37-3d | MATalpha his3Δ1 leu2Δ0 ura3Δ0 lys2Δ0 rpa34Δ::NATMX rpa49Δ::HPHMX RPA135-F301S::URA3 | This study |
| rpa14Δ in figure 4C | yCN225-1a | MATalpha his3Δ1 leu2Δ0 lysΔ0 ura3Δ0 rpa14Δ:: KANMX | This study |
| rpa14Δ RPA135-F301S in figure 4C | yTD38-3d | MATa his3Δ1 leu2Δ0 ura3Δ0 lys2Δ0 rpa14Δ::NATMX RPA135-F301S::URA3 | This study |
| rpa14Δ rpa49Δ RPA135-F301S in figure 4C | yTD39-8a | MATa his3Δ1 leu2Δ0 ura3Δ0 lys2Δ0 rpa14Δ::NATMX rpa49Δ::HPHMX RPA135-F301S::URA3 | This study |
| | yTD53-1a | MATalpha his3Δ1 leu2Δ0 lys2Δ0 ura3Δ0 KANMX6-pGAL::RPA12 | This study |
| RPA12 alleles in figure 5 with the plasmids indicated in the legend | OGT30-1c | MATa his3Δ1 leu2Δ0 lysΔ0 ura3Δ0 KANMX-pGAL::RPA12 | This study |
| rpa49Δ in figure 5 with the plasmids indicated in the legend | OGT30-3c | MATalpha his3Δ1 leu2Δ0 lysΔ0 ura3Δ0 rpa49Δ:: HPHMX KANMX-pGAL::RPA12 | This study |
| | OGT30-1a | MATa ura3-Δ0 his3-Δ1 leu2-Δ0 lys2-Δ0 rpa49Δ::HPH RPA135-(F301S)-TAP-HIS3 KANMX-pGAL::RPA12 | This study |
| | OGT30-3a | MATa ura3-Δ0 his3-Δ1 leu2-Δ0 lys2-Δ0 RPA135-(F301S)-TAP -HIS3 KANMX-pGAL::RPA12 | This study |
| | yTD40-1a | MATa ura3-Δ0 his3-Δ1 leu2-Δ0 lys2-Δ0 RPA135-(F301S)-STOP-URA3 KANMX-pGAL::RPA12 | This study |
| | yTD41-1a | MATa ura3-Δ0 his3-Δ1 leu2-Δ0 lys2-Δ0 rpa49Δ::HPH RPA135-(F301S)-STOP-URA3 KANMX-pGAL::RPA12 | This study |
| RPA135-F301S in figure 5C | yTD42-1a | MATa his3Δ1 leu2Δ0 lysΔ0 ura3Δ0 RPA135-F301S::URA3 KANMX-pGAL::RPA12 | This study |
| rpa49Δ RPA135-F301S in figure 5C | yTD43-1a | MATalpha his3Δ1 leu2Δ0 lysΔ0 ura3Δ0 rpa49Δ:: HPHMX RPA135-F301S::URA3 KANMX-pGAL::RPA12 | This study |
| rrp6Δ in figure 7 and S5 | yCD2-2a | MATa his3Δ1 leu2Δ0 ura3Δ met15Δ0 rrp6::NAT | This study |
| | yMKS8-1a | MATa his3Δ1 leu2Δ0 ura3Δ met15Δ0 rrp6::NAT KANMX6-pGAL::RPA49 | This study |
| Rpa135-F301S rrp6Δ in figure 7 and S5 | yMKS9-9d | MATa his3Δ1 leu2Δ0 ura3Δ rrp6::NAT RPA135-F301S-TAP-HIS3 | This study |

#### Supplementary table 3: Plasmids used in this study

| Name | Description | Source |
|---|---|---|
| pUC19-HPH | *Plasmid bearing HPH-MX4* | [55] |
| pFA6-kanMX6 | *Plasmid bearing KAN-MX6* | [56] |
| pFA6a-KanMX6-GAL::3HA | *Plasmid used for GAL promoter insertion* | [56] |
| pFA6a-HA-KlURA3 | *Plasmid used for epitope switching* | [57] |
| pNOY80 | *Plasmid CEN6 ARS4, URA3, RPA135* | [58] |
| pVV190 | *Plasmid pFL44-A190 (2µ URA3 RPA190)* | [24] |
| pRS316 | *CEN6 ARS4, URA3* | [59] |
| pCR4-HIS3 | *HIS3* | [59] [60] |
| pGL190_3 | *(2µ URA3 RPA190-E1274K) selected from randomly mutagenized p*VV190 | This study |
| pGL190_11 | *(2µ URA3 RPA190- C1493R) selected from randomly mutagenized p*VV190 | This study |
| pGL190_23 | *(2µ URA3 RPA190- L1262P) selected from randomly mutagenized p*VV190 | This study |
| pGL135_6prim | *(CEN4 URA3 RPA135-R379G selected from randomly mutagenized p*NOY80 | This study |
| pGL135_54 | *(CEN4 URA3 RPA135-Y252H) selected from randomly mutagenized p*NOY80 | This study |
| pGL135_33 | *(CEN4 URA3 RPA135-F301S) selected from randomly mutagenized p*NOY80 | This study |
| pTD1_3b_135TAP | *Plasmid pNOY80 bearing RPA135-TAP-HIS3 obtained by homologous recombination using a PCR-amplified fragment generated with oligos 835 and 836 and genomic DNA of strain 202233825 as template.* | This study |
| pTD2_6c_135TAP | *Plasmid pNOY80 bearing RPA135-F301S-TAP-HIS3 obtained by homologous recombination using a PCR-amplified fragment generated with oligos 835 and 836 and genomic DNA of strain 202233825 as template.* | This study |
| pTD5 | *Plasmid pTD1_3b_135TAP deleted of URA3 using NsiI and SdaI digestion and self-ligation.* | This study |
| pTD6 | *Plasmid pTD2_6c_135TAP deleted of URA3 using NsiI and SdaI digestion and self-ligation.* | This study |
| Ycp50-26 | *URA3 ARS/CEN RPA49* | [59] |
| pRS316-A12 | *pRS316 vector ligated with a PCR-generated fragment using oligos 1554 and 1555 and yeast genomic DNA as template cut BamHI-XbaI, and cloned at same site.* | This study |
| pRS316-A12-AvrII | *PCR mediated mutagenesis to introduce AvrII site in* pRS316-A12 using oligos 1714 and 1715. | This study |
| pRS316-A12-S6L | *S6L Allele of RPA12 isolated as a suppressor of the rpa49Δ growth defect, selected from a PCR-mediated random mutagenesis of* pRS316-A12-AvrII. | This study |
| pRS316-A12-S6L-KAN | *Plasmid obtained using* pRS316-A12-S6L *modified by homologous recombination using a PCR-amplified fragment generated with oligos 1682 and 1559 and* pFA6-kanMX6 *as template.* | This study |
| pRS316-A12-T49A | *T49A Allele of RPA12 isolated as a suppressor of the rpa49Δ growth defect, selected from a PCR mediated random mutagenesis of* pRS316-A12-AvrII. | This study |
| pRS316-A12-DCt | *Plasmid obtained using* pRS316-A12 *modified by homologous recombination using a PCR-amplified fragment generated with* oligos *1371 and 1559 and* pFA6-kanMX6 *as template.* | This study |
| pTD9 | *Plasmid obtained using* pRS316-A12-T49A *modified by homologous recombination using a PCR-amplified fragment generated with* oligos *1371 and 1559 and* pFA6-kanMX6 *as template.* | This study |
| pTD10 | *Plasmid obtained using* pRS316-A12-S6L *modified by homologous recombination using a PCR-amplified fragment generated with* oligos *1371 and 1559 and* pFA6-kanMX6 *as template.* | This study |
| pCJPF4 | *pFL36cII with LEU2 marker, CEN4, containing RPA49 coding region* | This study |
| pCJPF4-GAL49-1 | *Plasmid obtained using pCJPF4 modified by homologous recombination using a PCR-amplified fragment generated with* oligos *624 and 625 and* pFA6a-KanMX6-GAL::3HA *as template.* | This study |
| pMAX1 | *Plasmid including Pol I promoter used for in vitro assays* | [23] |
| pUC19tail_g-_601_elongated | *Plasmid used for tailed template* | [61] |

### Supplementary table 4: Oligonucleotides used in this study

| Sequence | N° |
|---|---|
| gctgctttatagaaccatctagcaaaaaaaaaaaaccagagcaaaccaaatgctcagtatatacctctaggtaactgaattcgagctcgtttaaac | 208 |
| aaagcagttgaagacaagttcgaa | 301 |
| gactctctccaccgtttgacg | 302 |
| ggtctgtgatgcccttagacg | 307 |
| agtttcacaagattaccaagacctctc | 308 |
| ggtggtaaattccatctaaagctaaatatt | 311 |
| cacgtactttttcactctcttttcaaa | 312 |
| ccactattgcagtcgttatcaaccttttgcactttatctagtagaattcgagctcgtttaaac | 624 |
| cactttcaatttcgatttcagaaacagaccttttcacggacattttgagatccgggtttt | 625 |
| catagtttctggcatgaccc | 649 |
| gaagaatcctggtagtatgg | 650 |
| tcgatgaattcgagctcgt | 700 |
| gtcttcaactgctttcgcat | 774 |
| gacgctgatgacatcgagtg | 835 |
| tgatgggcacatatgaagtc | 836 |
| catggcttaatctttgagac | 892 |
| aggttctaacgttaatgtttctcaa | 1189 |
| actttataataccctagctctctta | 1194 |
| aagtcacaattctccaatttaaaagtcgtcaccacgacggcagacgatgcgtttccatcttctcttagagccaagaaatgaggcgcgccacttctaaa | 1371 |
| ccatcgaaagttgatagggcag | 1501 |
| ctacgttcgacttatacaaaaaaaagaagtctgaaaaggacgaattcgtattacacggtgaaaacgagagactagaataccggatccccgggttaattaa | 1515 |
| gggggatcctttcaaatattgctataaaaatggatgatagc | 1554 |
| cgctctagacactgaatgtcacgatagagttatc | 1555 |
| cactgaatgtcacgatagagttatcgctgt | 1556 |
| tttcaaatattgctataaaaatggatgata | 1557 |
| tctatagatgttcacatgatgaaagcggggatgatattaatgtacaaattgtaatatgtgcgaacacaacccaatcagaattcgagctcgtttaaac | 1559 |
| ctaatacggcattaggattttccaggagatcaccacagtccaagcaaaaaattaacgatcctacaacagacattttgagatccgggtttt | 1634 |
| tttatgtcaagtgaagggcattgcatggcaagtgtaaagaatccaagaaggaattcgagctcgtttaaac | 1635 |
| agttaagatctgcagatgaaggtgctactgtcttctatacatgcacttcctgtggttacaagttccgtaccaacaattgatgaggcgcgccacttctaaa | 1682 |
| ttgaagtacttggactctgagctatccgcaatgggtataagattgcgttataatgtagagcccaaataacggatccccgggttaattaa | 1679 |
| gatctcctggaaaatcctaatgccgtcctaggctctaacgttgaatgcagcc | 1714 |
| ggctgcattcaacgttagagcctaggacggcattaggattttccaggagatc | 1715 |
| gatttgtttaacggcttctttgtagtcatcctgctgttgtttttgtatgtttaagcttcttcctccgattaagttttgtacatgtaaaacgacggccagt | 1716 |
| tgatgctttttgaagttttcatggcatgatttagcatttgaaatataatgagaaaagagccctttaacttacagaagtaaggaaacagctatgaccatg | 1717 |
| ttcttgcatgaacactaatgatattaggagggctttttaagttgctaccagaattcgagctcgtttaaac | 1711 |
| cactttcaatttcgatttcagaaacagaccttttcacggacatgcactgagcagcgtaatctg | 1713 |
| ctccgcttattgatatgc | 1829 |
| tgagaaggaaatgacgct | 1830 |
| accgtttggtctacccaagtgagaagccaagaca | 1831 |
| atcccggccgcctccatcac | 1832 |
| cggttttaattgtccta | 1833 |
| cctacagcgtgagctatgagaaag | 1834 |
| tcaccttaccctatacttactcg | 1835 |
| aaatggcctatcggaatacattttctacatcctaactactataaaacaacctttagacttacgtttgctactctcatggt | 1855 |
| tgcgaccggctattcaacaaggcattcccccaagtttgaattctttgaaatagattgctattagctagtaatccaccaaa | 1857 |
| ggaattcctcgttgaagagcaataattacaatgctctatccccagcacgacggagtttcacaagattaccaagacctctc | 1859 |
| gtgctggcctcttccagccataagaccccatctccggataaaccaattccggggtgataagctgttaagaagaaaagata | 1860 |
| gtaaatggtacactcttacacactatcatcctcatcgtatattataatagatatatacaatacatgtttttacccggatc | 1861 |
| tccgggcaaatcctttcacgctcgggaagctttgtgaaagcccttctctttcaacccatctttgcaacgaaaaaaaaaaa | 1862 |
| cagcttaactacagttgatcggacgggaaacggtgctttctggtagatatggccgcaaccgatagtttaacggaaacgca | 1863 |
| aaaaaaaaaaaagaaataaagattgcagcacctgagtttcgcgtatggtcacccactacactactcggtcaggctcttac | 1864 |
